# The subacromial bursa is a key regulator of the rotator cuff and a new therapeutic target for improving repair

**DOI:** 10.1101/2023.07.01.547347

**Authors:** Brittany P. Marshall, Xavier E. Ferrer, Jennifer A. Kunes, Astia C. Innis, Andrew J. Luzzi, Lynn Ann Forrester, Kevin G. Burt, Andy J. Lee, Lee Song, Clark T. Hung, William N. Levine, David Kovacevic, Stavros Thomopoulos

**Affiliations:** Department of Biomedical Engineering, Columbia University; New York, New York, USA; School of Medicine, Columbia University; New York, New York, USA; Department of Orthopedic Surgery, Columbia University; New York, New York, USA; New York Metropolitan Orthopaedics and Spine; New York NY, USA

## Abstract

Rotator cuff injuries result in over 500,000 surgeries performed annually, an alarmingly high number of which fail. These procedures typically involve repair of the injured tendon and removal of the subacromial bursa. However, recent identification of a resident population of mesenchymal stem cells and inflammatory responsiveness of the bursa to tendinopathy indicate an unexplored biological role of the bursa in the context of rotator cuff disease. Therefore, we aimed to understand the clinical relevance of bursa-tendon crosstalk, characterize the biologic role of the bursa within the shoulder, and test the therapeutic potential for targeting the bursa. Proteomic profiling of patient bursa and tendon samples demonstrated that the bursa is activated by tendon injury. Using a rat to model rotator cuff injury and repair, tenotomy-activated bursa protected the intact tendon adjacent to the injured tendon and maintained the morphology of the underlying bone. The bursa also promoted an early inflammatory response in the injured tendon, initiating key players in wound healing. *In vivo* results were supported by targeted organ culture studies of the bursa. To examine the potential to therapeutically target the bursa, dexamethasone was delivered to the bursa, prompting a shift in cellular signaling towards resolution of inflammation in the healing tendon. In conclusion, contrary to current clinical practice, the bursa should be retained to the greatest extent possible and provides a new therapeutically target for improving tendon healing outcomes.

**One Sentence Summary:** The subacromial bursa is activated by rotator cuff injury and regulates the paracrine environment of the shoulder to maintain the properties of the underlying tendon and bone.

## INTRODUCTION

Rotator cuff injuries cause pain, disability, and loss of shoulder function in over 17 million individuals in the United States, leading to over 500,000 surgeries performed annually with alarming failure rates of 20-94% (*1–4*). These surgeries typically involve repair or reconstruction of the damaged rotator cuff tendon(s) along with decompression of the subacromial space by debriding the overlying bone (acromion) and removing the subacromial bursa (*5–7*).

The subacromial bursa is a synovial-like tissue that is situated between the acromion and rotator cuff tendons. This tissue has long been described to serve a primarily mechanical role by providing cushioning and protection against friction-wear from the acromion on the underlying tendons. A pathological state in the bursa has been characterized by expression of inflammatory cytokines and markers of pain as well as tissue-level phenotypic changes (*8–10*). As a result of favorable improvements in pain scores following subacromial decompression, bursectomy is widely used to treat subacromial bursitis with or without a rotator cuff injury (*11–13*). Even in cases in which the bursa appears healthy intraoperatively, there is still clinical motivation for bursectomy because the bursa can obscure the surgical repair site (*8, 9, 14*). However, there is no evidence that decompression improves healing outcomes after rotator cuff repair (*5, 11, 13*). Bursectomy has even shown to be less effective in patients with subacromial pain if they also had a degenerative shoulder (*15*). Moreover, the identification of a robust vascular network within the bursa, a resident population of mesenchymal stem cells (MSCs), and inflammatory responsiveness to rotator cuff pathology have supported a biological role for this tissue in addition to its previously described mechanical role (*8, 9, 16–20*). These observations make surgical excision of the bursa problematic, given our current lack of understanding of the implications of removing the bursa on rotator cuff pathology and response to injury.

In addition to a putative biologic role, the subacromial bursa is also uniquely positioned superior to the rotator cuff, offering a biologically dynamic physical depot for drug delivery to the pathologic and/or healing rotator cuff. Further, the bursa is largely composed of adipose tissue (*20*), making it a natural carrier for lipophilic drug delivery systems. The bursa is therefore ideally suited as a therapeutic target to modulate shoulder inflammation, prevent rotator cuff degeneration, and enhance tendon-to-bone healing after surgical repair by promoting a paracrine environment supportive of regeneration. As current rehabilitation and surgical strategies have largely failed to prevent rotator cuff tears and improve healing after surgical repair, respectively, the bursa presents a transformative opportunity to improve outcomes in patients with rotator cuff pathologies.

Here, we *(i)* confirm the presence of bursa-tendon crosstalk in patients with rotator cuff pathology, *(ii)* establish that the bursa is activated by rotator cuff tenotomy in an animal model, *(iii)* identify an essential function of the bursa in regulating the paracrine environment in the shoulder, *(iv)* define the bursa’s role in driving the tendon’s healing response, and *(v)* demonstrate the therapeutic potential of targeting the subacromial bursa for modulating tendon inflammation.

## RESULTS

### Bursa-tendon crosstalk is present in patients with rotator cuff tears

In order to identify crosstalk between injured tendon and the overlying bursa in patients with tendinopathy, rotator cuff tendon and bursa tissue biopsies were analyzed using histologic and proteomics approaches. Intraoperative arthroscopic images of the bursa revealed three distinct subtypes: white fibrous (Fig. 1A), red vascular (Fig. 1B), and yellow fatty (Fig. 1C). Masson’s trichrome staining supported these gross observations, wherein the fibrous bursa stained primarily blue (connective tissue) (Fig. 1D), the vascular bursa had vascular elements (Fig. 1E), and the fatty bursa had vacuolated adipocytes (Fig. 1F). Analysis of the bursa proteome supported this tissue-level observation of distinct bursa subtypes, with the principal component analysis (PCA) clustering by subtypes (Fig. 1G). Prospective clinical data collection of study participants provided patient demographic and clinical metadata (Fig. S1 A-B), including sex, ethnicity, race, age, diagnoses, injury occurrence, and history of steroid injection. Using this clinical data, the bursa proteome was analyzed according to both bursa subtype (assigned on arthroscopic inspection) and tendon tear severity. The analysis of the bursa proteome by tendon tear severity indicated clustering in the heat map and PCA defined by the presence of a tear (Fig. 1G-H). When analyzed for the contribution of age to proteome clustering, one outlier in age was explained by a prior revision surgery (Fig. S1 C). Furthermore, one outlier within the fibrous subtype was explained by age, as it was the one fibrous sample belonging to a subject over 60 years old (Fig. S1 C). These results suggest that the bursa proteome is defined by a combination of the factors: subtype, age, and tendon tear presence. Proteomic analysis of the tendon indicated clustering of the tendon proteome by tear size within the PCA (Fig. S1 D). Hierarchical heat map analysis supported the results from the PCA (Fig. S1E), indicating that injury to the tendon drives a molecular response in the adjacent bursa.

**Fig. 1:**
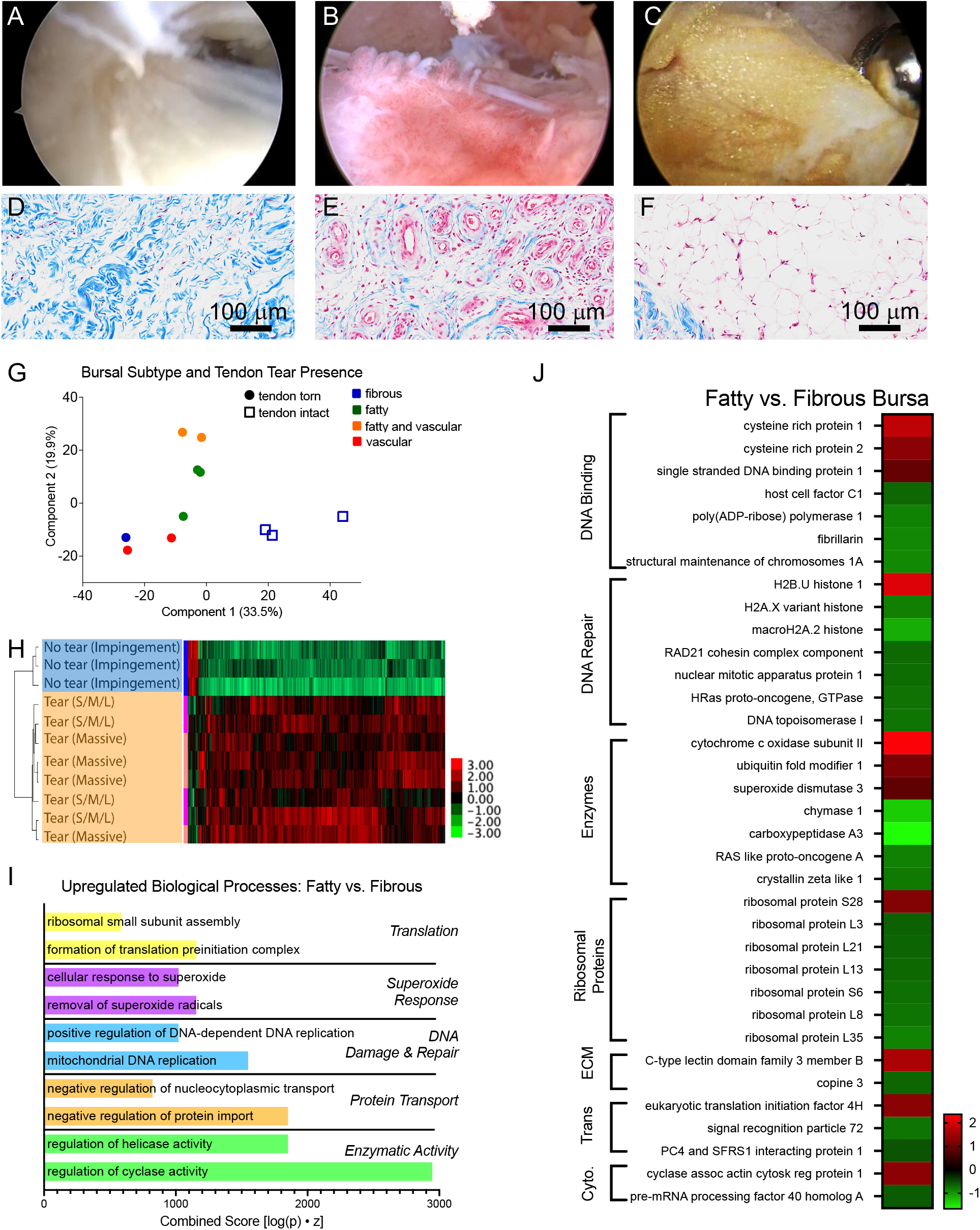
The bursa is activated in patients with rotator cuff tendinopathy. (A-C) Arthroscopic images and (D-F) Masson’s trichrome histological staining of the bursa, identified as (A, D) fibrous, (B, E) vascular, or (C, F) fatty subtypes. (G) Principal component analysis of bursa biopsies indicated separation of the bursa proteome by subtype and by presence of a tear in the associated tendon. (H) Hierarchical clustering heatmap indicated clear separation of the bursa proteome by tendon tear presence. (I) Pathway analysis revealed biological processes upregulated in the fatty compared to fibrous bursa subtypes, with (J) specific differentially regulated genes implicated in these pathways.

Further investigation into the specific protein-level changes in the vascular, fatty, and fibrous bursa subtypes revealed unique protein signatures and one shared protein (SOD3) in the vascular compared to fibrous and fatty compared to fibrous bursa analyses (Fig. S1 F). Gene ontology process analysis of the fatty vs. fibrous comparison indicated differences in translation, superoxide response, DNA damage and repair, protein transport, and enzymatic activity processes (Fig. 1I). These processes were in contrast to those within the vascular vs. fibrous comparison, which included cell movement, superoxide response, complement cascade, cytokine and immune cell signaling, fatty metabolism, and extracellular matrix related pathways (Fig. S1 G). Heatmaps of specific genes supported these process-level findings in the fatty vs. fibrous bursa, indicating proteins with roles in DNA binding and repair, enzymatic processes, ribosomes, extracellular matrix, transcription and translation (trans), and the cytoskeleton (cyto) (Fig. 1J). In contrast, proteins in the vascular bursa had roles in extracellular matrix, vascular, enzymatic, immune response, lipid metabolism, and mechanoreceptive functions (Fig. S1 H).

### The bursa is activated by tendinopathy

The evidence of clinically relevant crosstalk between the bursa and the rotator cuff motivated focused investigation of bursa-tendon crosstalk using two models of tendinopathy (one *in vivo* and one *in vitro*) (Fig. 2A). The rat was first validated as a clinically relevant model for investigation of bursa-rotator cuff tendon crosstalk, including consideration of the anatomic location of the bursa and the presence of resident MSCs (as previously demonstrated in human bursa studies (*19*)). Dissection of the rat shoulder revealed a glistening pink/yellow bursa located inferior to the trapezius muscle in its medial portion, inferior to the acromion in its distal portion, and superior to the supraspinatus (Fig. S2 A-B). Microcomputed tomography reconstructions of contrast-enhanced whole-shoulder scans revealed a region of low radiopacity inferior to the acromion and trapezius and superior to the supraspinatus tendon (Fig. S2 C). Histological assessment of the rat shoulder (sectioned in the coronal plane) and stained with Movat’s pentachrome supported this finding, revealing an adipose-rich subacromial bursa inferior to the trapezius muscle and acromion and superior to the supraspinatus tendon of the rotator cuff (Fig. 2B).

**Fig. 2:**
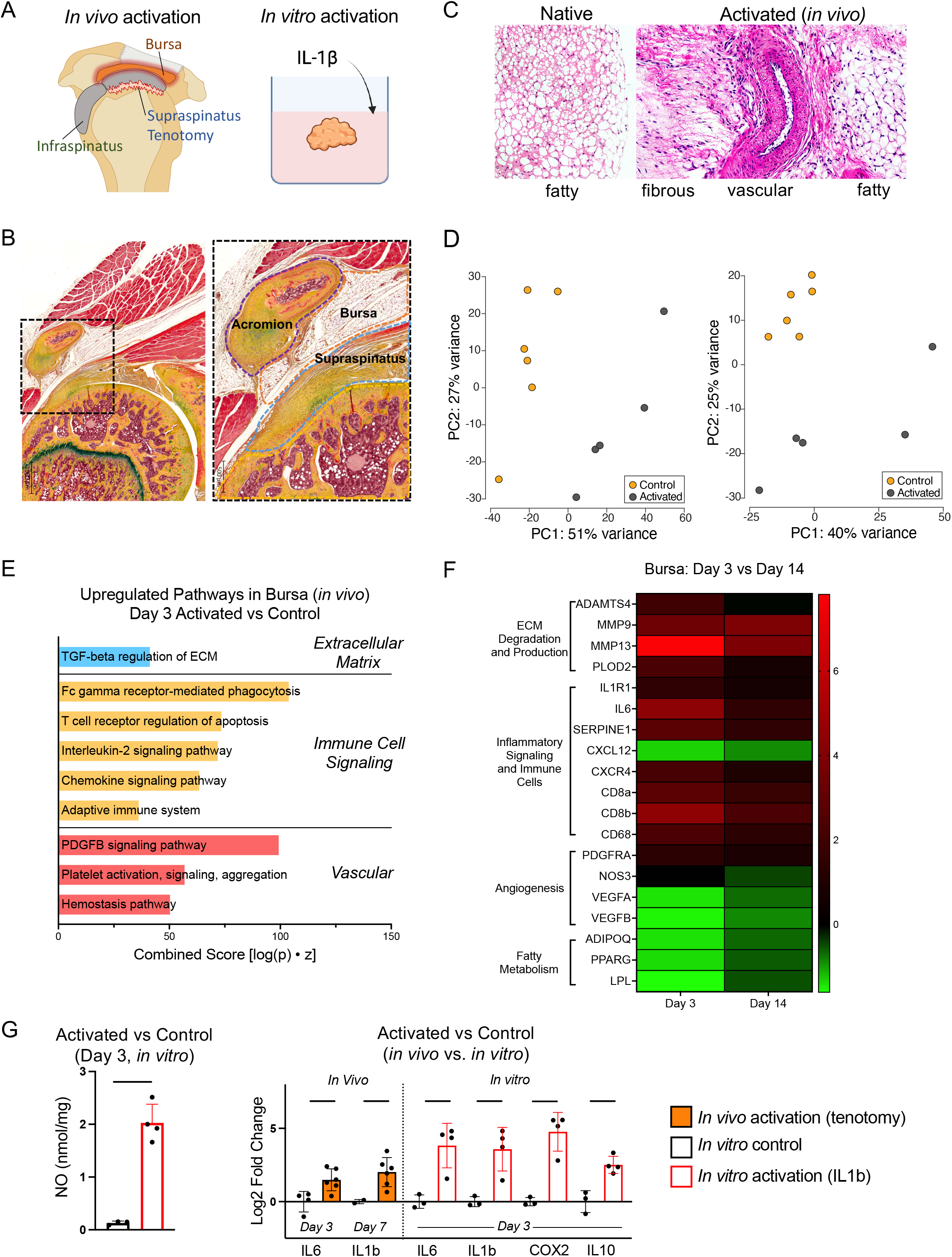
The bursa is activated by the inflammatory milieu present in models of tendinopathy. (A) Bursa activation was studied in vivo with a tenotomy and in vitro with the addition of proinflammatory cytokine IL1β. (B) Movat’s pentachrome-stained whole shoulder histological section revealed the subacromial bursa positioned between the acromion and supraspinatus tendon. (C) H&E staining showed bursa activation, induced by tenotomy, led to a transition from fatty in its native state to fibrous, vascular, and fatty in its activated state. (D) PCA at days 3 and 14 indicated clear separation of activated from control bursa transcriptomes. (E) Pathway analysis of day 3 bursa revealed specific pathways upregulated in the activated bursa, consistent with (F) specific genes that play key roles in these pathways; the magnitude of differential expression was higher at day 3 than at day 14. (G) Comparison of in vivo (orange) and in vitro (red outline) bursa activation revealed similar gene expression patterns in both models, with NO release also increased in the in vitro activation. [p < 0.05 indicated by line over bars]

To confirm the presence of MSCs within the rat bursa, adherent cells from the bursa were isolated and cultured. Using flow cytometry and immunocytochemistry, these bursa-derived cells were confirmed to have known markers for MSCs, including CD29+, CD73+, CD90+, and CD105+, and were also CD45-(Fig. S3 A-D). Pluripotency of this cell population was also confirmed with multilineage differentiations into chondrogenic, adipogenic, osteogenic, and tenogenic lineages. Adipogenic and osteogenic differentiation cultures were stained and confirmed positive for lipid deposition (Fig. S3 E) and mineral deposition (Fig. S3 F), respectively. Furthermore, gene expression confirmed successful differentiation towards chondrogenic (ACAN, COL2a1, SOX9), adipogenic (LPL, ADIPOQ, PPARγ), osteogenic (ALP), and tenogenic (SCX, TNMD) lineages due to the respective differentiation conditions (Fig. S3 G). Together, these assays confirmed that the rat bursa, like the human bursa, contains a resident population of MSCs.

The rat model was then used to determine the presence of bursa-tendon crosstalk during tendinopathy, which was modeled *in vivo* by a supraspinatus tenotomy (Fig. 2A). The bursa was activated to a state of cellular infiltration and heightened transcription by the nearby tendon injury, as demonstrated by histology, transcriptomics, and gene expression (Fig. 2C-G). Given 56 days of healing following bursa activation, phenotypic shifts towards fatty, vascular, and fibrous bursal subtypes were observed histologically in a manner consistent with the clinical bursa histology (Fig. 2C). Transcriptomic analysis of the activated bursa indicated unique and overlapping genes between days 3 and 14 post-tenotomy that were differentially regulated compared to control (Fig. S4 A-B). PCA demonstrated clear separation of the activated bursa from the control bursa at both timepoints (Fig. 2D). Building on the cellular characterization of MSCs in the healthy rat bursa, cell-type analysis of the day 3 activated bursa transcriptomics revealed several upregulated cells, including immune cells (i.e., monocytes, dendritic cells, B-cells, natural killer cells) and vascular cells (i.e., endothelial cells and early erythroid) (Fig. S4 C). Similarly, at day 14 the activated bursa transcriptomics indicated upregulated immune cells (i.e., monocytes, dendritic cells, natural killer cells, CD8+ and CD4+ T-cells, and B-cells) and vascular cells (i.e., early erythroid, and CD105+ endothelial cells) (Fig. S4 C). Pathway analysis of the bursa transcriptomics data indicated upregulated ECM, immune-cell signaling, and vascular-related pathways at day 3 (Fig. 2E), and upregulated ECM, innervation, immune cell signaling, and cytokine signaling pathways at day 14 in the activated bursa compared to control (Fig. S4 D). An investigation into the magnitude of up- or down-regulation of key genes within ECM-, inflammatory-, angiogenic-, and fatty metabolism-related pathways indicated a greater magnitude of activation in the bursa at day 3 compared to day 14 (Fig. 2F). This result was support by individual gene expression, which showed upregulation of inflammatory genes in the bursa, including IL6 at day 3 and IL6, IL1β, and TNFα at day 7 post-injury; genes that were no longer upregulated by day 14 (Fig. S4 E).

Using an *in vitro* organ culture model, the bursa was similarly activated by IL1β, a key cytokine of the inflammatory milieu resulting from tendon injury (Fig. 2A) (*21*). This state of inflammatory activation was indicated by increased levels of nitric oxide (NO) compared to control (Fig. 2G). Correspondingly, the IL1β-activated bursa upregulated expression of IL6, IL1b, COX2, and IL10 (Fig. 2G), similar to the *in vivo* animal model. Importantly, explanted bursa tissue viability was confirmed with a live (green)/dead (red) stain at 8 days in complete media (Fig. S5 A) and at 72 hours when activated with IL1β (Fig. S5 B), compared to a kill control (Fig. S5 C).

### The activated bursa regulates its paracrine environment to maintain the health of the underlying tendon and bone

Having established the bursa’s responsiveness to the inflammatory milieu produced during tendinopathy, the activated bursa’s role in regulating the paracrine environment within the shoulder, including the intact infraspinatus tendon and the underlying humeral head bone, was investigated. Using the *in vivo* model of tendinopathy (induced via supraspinatus tenotomy), a paired study design was used (bursectomy was performed unilaterally) to examine the bursa’s role for maintaining the health of intact tissues of the shoulder via paracrine regulation (Fig. 3A).

**Fig. 3:**
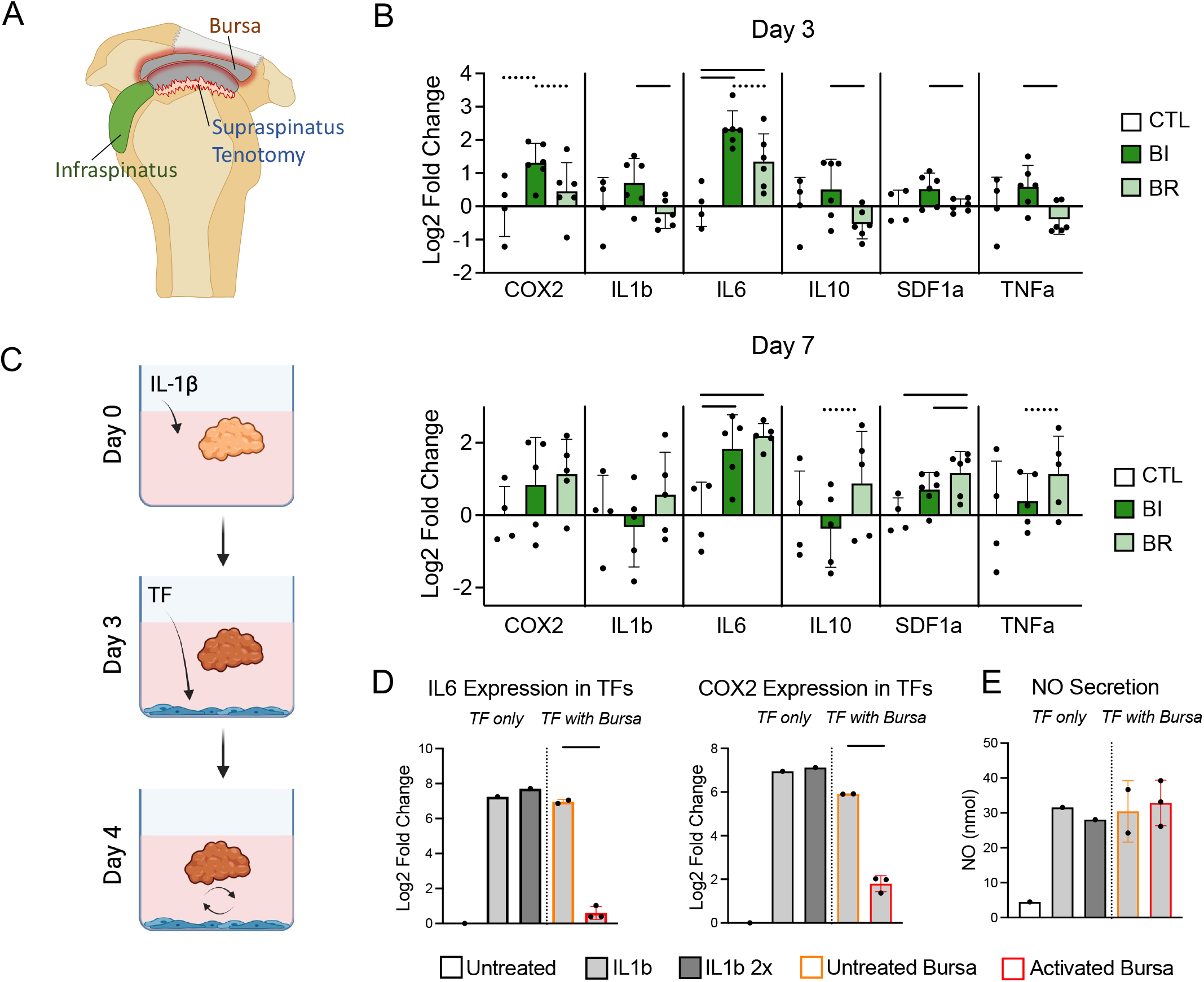
The activated bursa regulates its paracrine environment to maintain the health of the underlying infraspinatus tendon. (A) The role of the bursa in regulating the inflammatory environment in the shoulder was investigated within the intact rotator cuff (infraspinatus tendon) following supraspinatus tenotomy. (B) At day 3, expression of inflammatory mediators was upregulated by the bursa, but by day 7 the bursa drove down this inflammatory expression. (C) Resolution of inflammation by the activated bursa was investigated in vitro with IL1β activation of the bursa and tendon fibroblasts before co-culture. (D) Expression of IL6 and COX2 in the tendon fibroblasts decreased in co-culture with the activated bursa. (E) NO secretion was elevated in all conditions except untreated control tendon fibroblasts. [p < 0.05 indicated by line over bars; p<0.01 indicated by dotted lines; CTL: control, BI: bursa intact, BR: bursa removed; IL1b 2x indicated repeat dose of IL1b].

In the setting of a supraspinatus injury, removal of the bursa was detrimental to the inflammatory environment and biomechanical properties of the infraspinatus tendon and the bone morphometric properties in the humeral head. The bursa’s role in the intact infraspinatus response to supraspinatus injury was time-dependent. At day 3, the bursa drove up the inflammatory response in the infraspinatus, as indicated by increased expression of IL1β, IL10, SDF1a, TNFα, COX2 (trend), and IL6 (trend) (Fig. 3B). At day 7, this behavior reversed, and the bursa resolved inflammatory expression, including for SDF1a, IL10 (trend), and TNFα (trend), suggesting an immunomodulatory role of the bursa (Fig. 3B). By day 14 post-intervention, the bursa had little effect on expression of the set of genes screened at days 3 and 7 (Fig. S6 A). Similarly, using the *in vitro* model of crosstalk between tendon fibroblasts and the bursa (Fig. 3C), gene expression analysis of tendon fibroblasts revealed a clear immunomodulatory role by the activated bursa. This immunomodulatory role was evidenced by decreased COX2 and IL6 expression in the IL1β pre-treated tendon fibroblasts co-cultured with activated bursa relative to control bursa (Fig. 3D). NO secretion into the media for all treatment conditions except untreated tendon fibroblasts were similarly elevated (Fig. 3E). Decreased expression of IL6 and COX2 and increased NO secretion was seen when co-cultured with activated bursa using an *in vitro* model with untreated tendon fibroblasts (Fig. S6 B-C)). Together, these data, representing resident and infiltrating cells *in vivo* and isolated resident cells *in vitro* tell a compelling story of the activated bursa’s capacity for immunomodulation via crosstalk with tendon. The results demonstrate clear involvement of the bursa in the inflammatory response to injury in the shoulder.

To examine longer-term consequences of bursectomy in the context of tendinopathy, the uninjured infraspinatus tendon was studied at days 14, 28, 56 post-supraspinatus injury and repair. The bursa played a time-dependent role in the longer-term health of the intact infraspinatus tendon in the environment of the repaired (healing) supraspinatus tendon (Fig. 4A). Mechanical testing of the infraspinatus demonstrated an important role of the bursa in maintaining native properties of the intact rotator cuff. Specifically, based on quasistatic tensile tests to failure, the strength (i.e., maximum force) and stiffness (i.e., slope of the load-deformation curve) of the infraspinatus in the setting of a healing supraspinatus tendon were equivalent to control tendons when the bursa was intact, and significantly higher than infraspinatus tendons in the bursectomy group (Fig. 4B). Consistent with these mechanical results, the bursa suppressed early ECM-related gene expression (day 14 – COL3, BGN, COL1 (trend)), but enhanced later ECM-related gene expression (day 56 – COL1, COL3) in the infraspinatus tendon (Fig. 4C). The bursa also promoted day 14 expression of the tenogenic gene SCX (Fig. 4C). In the interim between these timepoints (day 28), the bursa appeared to have little effect on ECM-related gene expression in the infraspinatus (Fig. S6 D). Increased collagen expression (the primary load-bearing component of tendon ECM) in the later phase of healing is consistent with the beneficial role of the bursa on the mechanical properties of the infraspinatus tendon.

**Fig. 4:**
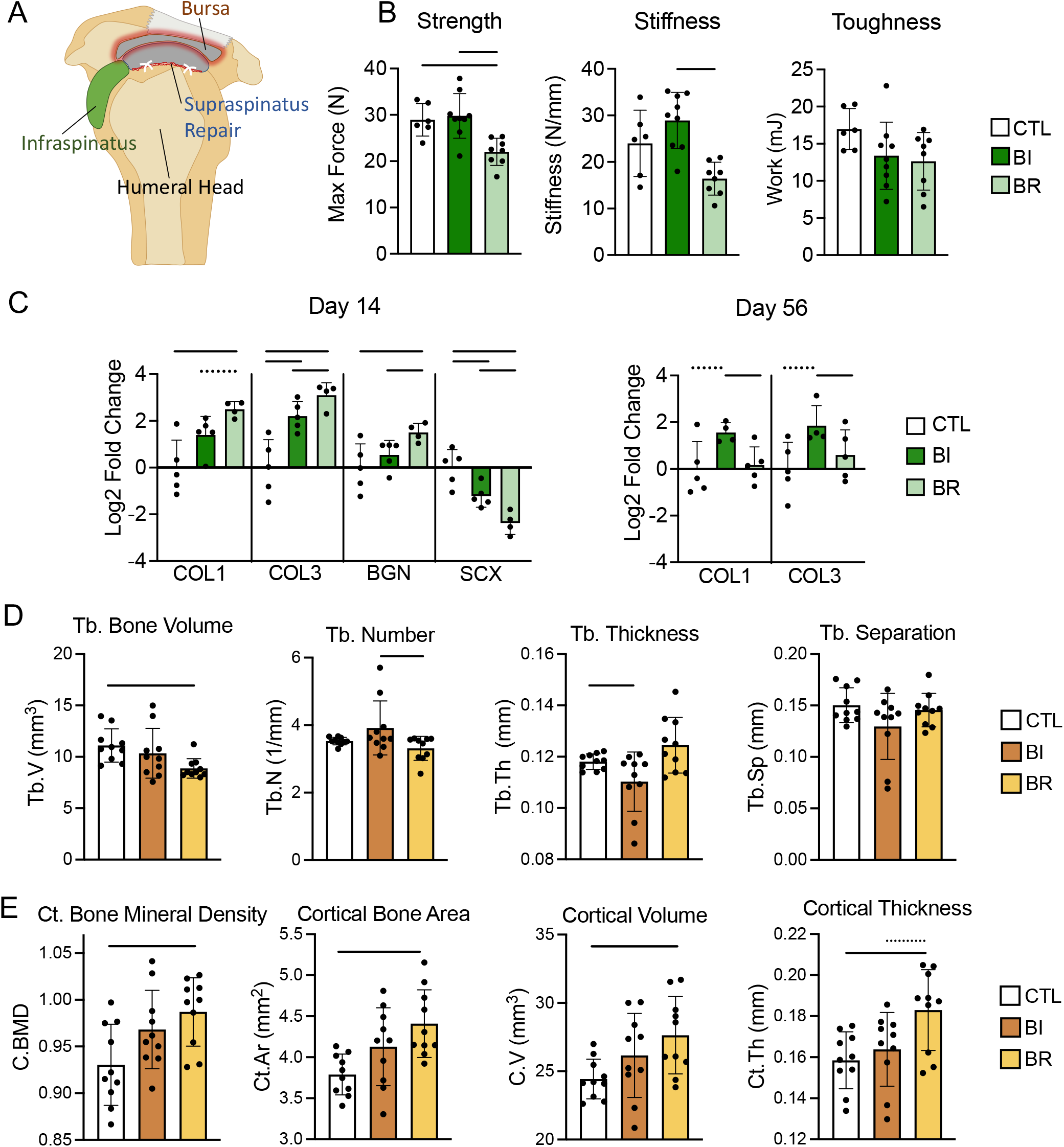
The activated bursa regulates its paracrine environment to maintain the health of the underlying humeral head bone. (A) The role of the bursa in regulating the healing environment in the shoulder was investigated within the intact rotator cuff (infraspinatus) and underlying humerus following supraspinatus tenotomy and repair. (B) Mechanical testing of the infraspinatus tendon revealed a decrease in strength and stiffness when the bursa was removed. (C) Infraspinatus tendon gene expression for ECM-related genes and the tenogenic transcription factor SCX were bursa-dependent. (D) Humeral head trabecular bone morphometry revealed decreased trabecular bone volume and number with the bursa removed. Trabecular thickness was decreased with the bursa intact. There was no change in trabecular separation due to the bursa. (E) Cortical bone morphometry indicated maintenance of cortical bone mineral density, area, volume, and thickness with the bursa intact and increases in all four parameters with the bursa removed, compared to control. [p < 0.05 indicated by line over bars, p < 0.1 indicated by dotted line over bars; CTL: control, BI: bursa intact, BR: bursa removed].

The results for the infraspinatus tendon indicated that the bursa supports a favorable paracrine environment in the shoulder. Therefore, we next examined the morphology of the humeral bone underlying the rotator cuff, which typically loses bone during tendon-to-bone healing. The loading environment of the humeral head, which plays a critical role in maintaining bone morphology, was identical in both bursa intact and bursa removed cases. Consistent with the results in the intact infraspinatus tendon, trabecular and cortical bone properties in the humeral head were dependent on the presence of the bursa during healing. Specifically, trabecular bone volume and trabecular number were decreased when the bursa was removed (Fig. 4D). Cortical bone mineral density, area, volume, and thickness increased when the bursa was removed (Fig. 4E). These results indicate deviations from native bone morphometry that were unique to the bursectomy group.

### The bursa directs the healing response in tendinopathy

The *in vivo* and *in vitro* results above demonstrated a clear beneficial role of the activated bursa in regulating the paracrine environment within the shoulder following tendon injury, specifically regarding the uninjured infraspinatus tendon and the humeral head bone. To investigate the role of the bursa on the injured tendon itself, the repaired supraspinatus tendons from the acute injury model were analyzed for transcriptional, histological, and biomechanical changes with and without bursectomy (Fig. 5A).

**Fig. 5:**
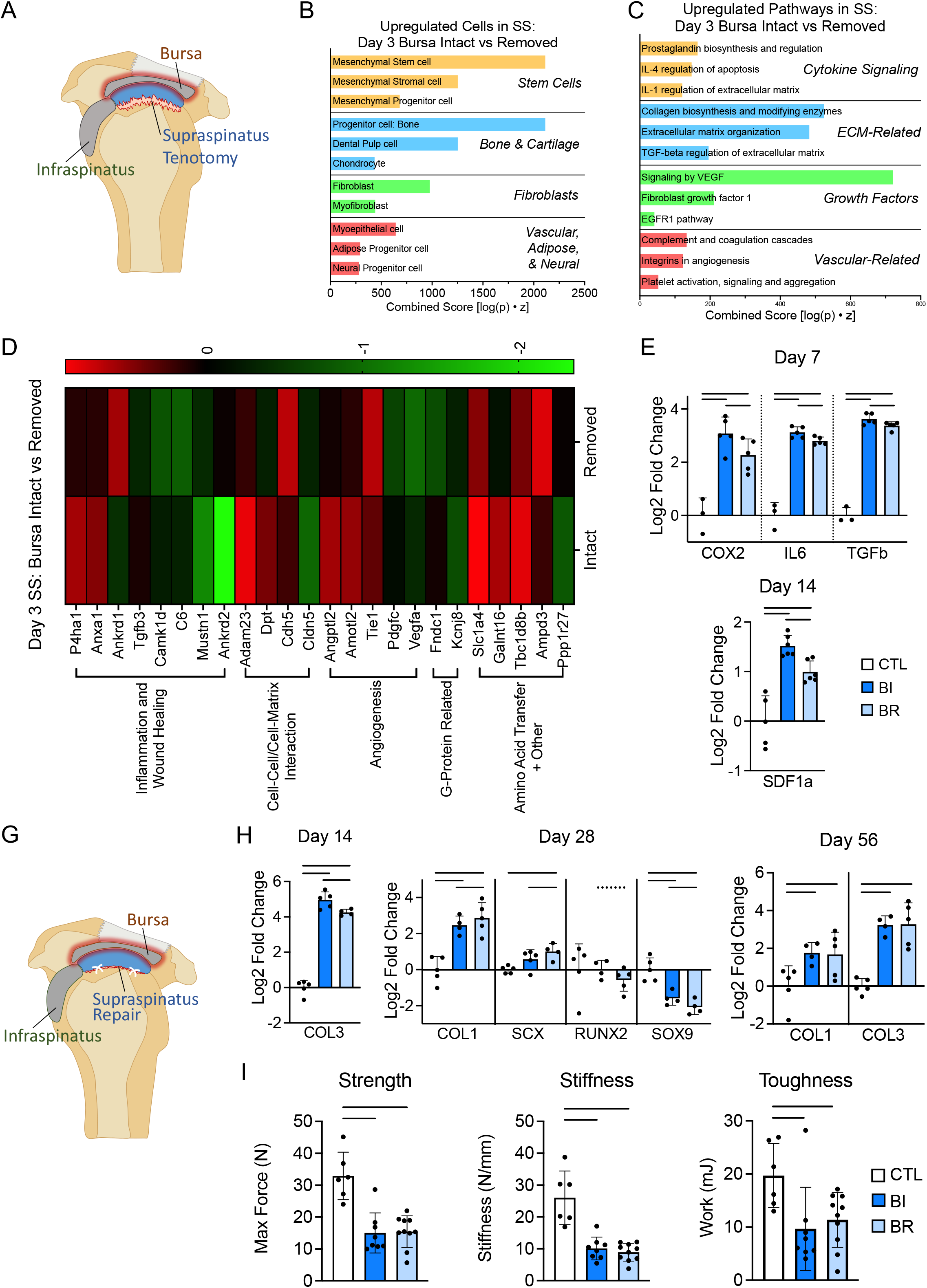
The bursa regulates the inflammatory, proliferative, and remodeling stages of tendon healing. (A) The role of the bursa in regulating the tendon inflammatory response to injury was investigated in the supraspinatus tendon following tenotomy. (B) Transcriptomic cell-type analysis at day 3 revealed multiple cell types upregulated in the injured tendon by the bursa. (C) Pathway analysis revealed several types of pathways upregulated by the bursa that were supported by (D) heat map of select genes up- and down-regulated in the supraspinatus tendon by the bursa. (E) The bursa upregulated expression of inflammatory markers COX2, IL6, and TGFβ at day 7 and stem- and immune cell homing factor SDF1a at day 14. (G) The role of the bursa in regulating tendon healing was investigated within the supraspinatus following tenotomy and repair. (H) The bursa increased expression of COL3 at day 14, decreased expression of COL1, SCX, and RUNX2 (trend) and increased expression of SOX9 at day 28, and had no effect on COL1 or COL3 expression by day 56. (I) These ECM-related gene expression changes during the early healing stages resulted in no difference in strength, stiffness, or toughness in the healing tendon by day 56. [p < 0.05 indicated by line over bars, p < 0.1 indicated by dotted line over bars; CTL: control, BI: bursa intact, BR: bursa removed].

Transcriptomic analysis of the supraspinatus tendon at day 3 after injury indicated both unique and shared up- and down-regulated genes between the bursa intact and the bursa removed groups (Fig. S7 A). A cell-type analysis of the transcriptome indicated that the bursa upregulated gene sets associated with stem cells, bone and cartilage cells, fibroblasts, and vascular, adipose, and neural cells (Fig. 5B). Pathway analysis showed that cytokine signaling, ECM, growth factors, and vascular-response pathways were upregulated by the bursa (Fig. 5C). A heat map of those genes differentially expressed due to bursectomy indicated that genes related to inflammation and wound healing, cell-cell or cell-matrix interaction, angiogenesis, G-proteins, and amino acid transfer were differentially expressed depending on the presence of the bursa (Fig. 5D). When examining specific genes with RT-qPCR, the bursa drove up inflammatory gene expression (day 3-COX2; day 7-COX2, IL6, TGFβ; day 14-SDF1a) in the injured supraspinatus tendon, supporting the transcriptomic evidence of upregulated cytokine signaling pathways in the bursa intact group (Fig. S7 B, Fig. 5E). At the tissue level, histologic analysis showed that the bursa imparted a trend towards increased cellular infiltration into the injured tendon by day 14 (Fig. S7 C). Together, these results point to a robust involvement of the bursa in directing cellular activity during the early period after tendon injury.

To understand the longer-term role of the bursa on tendon healing, we examined tendon healing 14 through 56 days after injury and repair. Bilateral supraspinatus tendon injuries were created and immediately repaired, and unilateral bursectomies were performed to test the role of the bursa on healing (Fig. 5G). The bursa drove down the expression of tenogenic gene SCX and tendon ECM gene COL1 at day 28 but drove up the expression of scar-mediated healing gene COL3 at day 14 and chondro/osteo-genic genes SOX9 and RUNX2 at day 28 in the injured and repaired supraspinatus (Fig. 5H). The bursa had no effect on COL1 or COL3 expression by day 56 (Fig. 5H). Looking to the functional consequences of these transcriptional changes, the bursa imparted no effect on the strength, stiffness, and toughness of the supraspinatus tendon when pulled to failure in uniaxial tension (Fig. 5I).

### The bursa is a therapeutic target for improving tendon healing

It is well established that rotator cuff repairs have high failure rates due to poor tendon-to-bone healing. The above data demonstrated that the bursa beneficially regulates the paracrine environment of the shoulder, supporting the recommendation against bursectomy during rotator cuff repair/reconstruction. Furthermore, the bursa, with its proximity to the rotator cuff, cellular responsivity, and ECM capable of immobilizing a lipophilic drug, is well-positioned for *in situ* drug delivery for improving rotator cuff healing. Corticosteroids such as dexamethasone (Dex), which have the capacity to dampen cytokine-mediated inflammation and have been previously used to treat inflammatory musculoskeletal pathologies, are candidate drugs for bursa-targeted rotator cuff treatments.

The efficacy of Dex delivery to the bursa for dampening its activated state was investigated *in vitro* by pre-treating bursa explants with IL1β prior to adding solubilized Dex to the culture media (Fig. 6A). The addition of Dex decreased expression of IL6 and IL1β and NO secretion in the activated bursa (Fig. 6A). Bursa activated with IL1β and treated with Dex were then co-cultured with IL1β pre-treated tendon fibroblasts (Fig. 6B). Gene expression of the tendon fibroblasts after 24 hours of bursa-tendon crosstalk revealed that Dex further enhanced the immunomodulatory capacity of the activated bursa. This was demonstrated by decreased IL6 expression in the tendon fibroblasts cultured with the IL1β- and Dex-conditioned bursa compared to the co-culture with the IL1β-activated or control bursa (Fig. 6B). The addition of Dex to the IL1β-activated bursa also decreased NO secretion into the media compared to the co-culture with IL1β-activated bursa (Fig. 6B). These findings supported the use of Dex for modulating the cytokine-mediated inflammatory response in the bursa and, in turn, modulating the inflammatory response in the tendon.

**Fig. 6:**
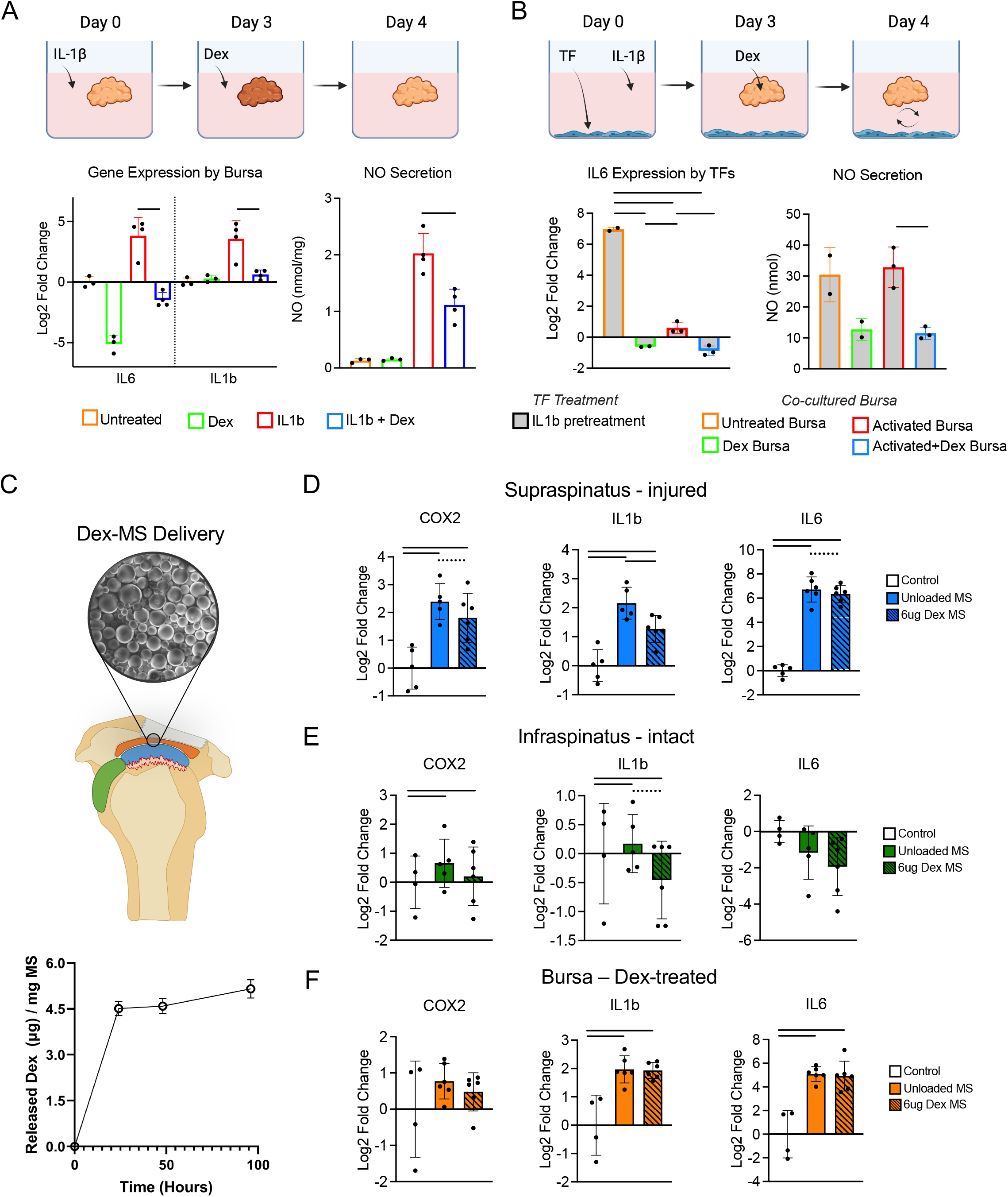
Dexamethasone delivery to the bursa regulates the tendinopathic inflammatory response. (A) The efficacy of Dex for decreasing IL1β-induced activation in the bursa was investigated in vitro. IL6 and IL1β expression and NO release were decreased with the addition of Dex. (B) The activated and Dex-treated bursa were co-cultured with IL1β-pre-treated tendon fibroblasts to screen Dex as a therapeutic candidate for reducing tendinopathic inflammation. The Dex-treated, activated, and activated and Dex-treated bursa decreased expression of IL6 in the co-cultured tendon fibroblasts. The addition of Dex to the activated bursa also decreased NO secretion in co-culture compared to the activated bursa. (C) Bursa-targeted Dex delivery was translated for in vivo use by encapsulation in 10-80 μm hydrophobic/lipophilic PLGA microspheres that release Dex in a burst over 24 hours followed by a slow release through 96 hours. (D) Dex-MS delivery to the bursa decreased IL1β, COX2 (p=0.10), and IL6 (p=0.089) expression in the injured supraspinatus tendon. (E) Dex-MS delivery to the bursa decreased IL1β (p=0.060) expression in the intact infraspinatus tendon. (F) Dex-MS delivery to the bursa had no effect on expression of COX2, IL1β, or IL6 in the bursa itself. [p < 0.05 indicated by line over bars, p < 0.1 indicated by dotted line over bars].

Translation of this therapeutic approach *in vivo* required a delivery system for sustained release of Dex, as repeat dosing is not practical in a clinical setting. Further, high doses of solubilized corticosteroid are required due to high clearance rates in joint tissues that result in short residence times with high potential local toxicity. Therefore, Dex was encapsulated in 10-80 µm PLGA microspheres (MS), as validated by scanning electron microscopy (Fig. 6C). Quantification of Dex release from the Dex-MS into phosphate buffered saline (PBS) indicated a burst release in the first 24 hours followed by slow release through 96 hours (the longest timepoint assessed *in vivo*) (Fig. 6C). Dex-MS with 6µg Dex were suspended in a 3µL PBS bolus and injected into the bursa *in vivo* immediately after supraspinatus injury; gene expression and histology were assessed 3 days later. All results were compared to a contralateral supraspinatus injury with injection of unloaded MS into the bursa. The Dex-MS decreased IL1β expression and led to a trend towards decreasing COX2 and IL6 expression in the injured supraspinatus tendon (Fig. 6D). Dex-MS also led to a trending decrease in IL1β expression in the intact infraspinatus tendon (Fig. 6E). Dex-MS did not change expression of COX2, IL1β, or IL6 in the bursa itself, although only a limited number of candidate genes were examined; transcriptomics/proteomics is necessary to identify potential drivers of the bursa-initiated changes seen in the rotator cuff tendons (Fig. 6F).

Clinically, most rotator cuff tears do not present for surgical intervention immediately after injury, in contrast to the animal model where injury, repair, and drug delivery were performed concurrently. For most patients, there is a delay between the onset of injury, assessment of the injury by a physician, and initiation of a treatment. To better model this clinical scenario, we investigated the efficacy of delayed delivery of Dex-MS (6µg Dex; 3µL PBS) to the bursa 3 days following the supraspinatus tendon injury (Fig. S8 A). Here, there was a trend towards a decrease in COX2 and an increase in SDF1a expression in the supraspinatus tendon (Fig. S8 B). Similarly, there was an increase in SDF1a expression in the infraspinatus tendon (Fig. S8 C). Consistent with the findings of the study where treatment was applied concurrently with injury and repair, no change in the expression of COX2, IL1 β, IL6, or SDF1a was observed in the delayed Dex-treated bursa itself compared to control (Fig. S8 D). Together, these studies serve as preliminary evidence in support of bursa-targeted drug delivery for modulating rotator cuff tendon responses to injury.

## DISCUSSION

Motivated by evidence from patient biopsies, a critical role of the subacromial bursa was demonstrated for rotator cuff pathology and healing in *in vitro* and *in vivo* model systems. Sampling of patient tissues revealed distinct proteomic signatures of the bursa and rotator cuff tendon that depended on bursa tissue subtype and tendon tear severity, respectively. Associations between the bursa proteome and the underlying tendon tear provided compelling evidence of crosstalk between the two tissues. These findings are consistent with a recent study that found upregulated markers of oxidative stress and chronic inflammation in bursa from patients with rotator cuff tears compared to those with shoulder instability, indicating that the bursa proteome was dependent on the pathological state of the rotator cuff (*22*). To control the state of the bursa and tendon and test for causal crosstalk links between the tissues, bursa organ cultures and a rat model of surgical injury and repair were used. These models permitted *in vitro* and *in vivo* investigation of the bursa in its healthy and pathological states and allowed us to pose questions that cannot be answered with pathologic patient samples. Activation of the bursa by tenotomy of the underlying rotator cuff tendon led to infiltration of the bursa by an array of immune cells and phenotypic shifts at the tissue and molecular levels, consistent with previous observations of the pathological human bursa. Santavirta et al determined that the pathological bursa possessed 50-80% CD2+ T-cells, 10-40% CD11b+ macrophages, and very few PCA1+ plasma cells (*23*). A more recent study demonstrated the presence of CD68+ macrophages in all bursa samples examined from patients with shoulder instability, osteochondral pathology, or partial-to-full thickness supraspinatus tears (*10*). These findings are consistent with our transcriptomics results from 3 and 14 day activated bursa, which both contained immune cells, including T-cells, monocytes, and B-cells. Similarly, a number of studies have shown that the pathological human bursa has upregulated expression of IL1α, IL1β, IL6, TNFα, MMPs, COX1/2, and VEGF (*8, 9, 24, 25*), including a study that correlated increased expression of IL1β, IL6, and COX2 in the bursa with increasing severity of supraspinatus tear (*10*). Gene expression from the activated rat bursa demonstrated upregulated IL1β, IL6, and TNFα, and transcriptomics demonstrated upregulated immune cell and cytokine signaling pathways. These cellular and gene expression results together demonstrate that the bursa is responsive to the inflammatory environment generated during both rat and human tendinopathy.

Surgical removal of the bursa has long been motivated by the assumption that it adversely contributes to inflammation and catabolism in the shoulder. However, the current work demonstrates that the activated bursa can play a beneficial role, particularly for resolving inflammation and protecting the intact infraspinatus tendon adjacent to the injured supraspinatus tendon. Importantly, this immunomodulatory capacity was only observed when the bursa was activated by the inflammatory milieu of an injury setting, and demonstrated most clearly in the *in vitro* model of bursa-tendon crosstalk. The bursa both incites and resolves inflammation in the surrounding tissues, allowing for a more rapid transition into the proliferative and remodeling phases of healing. The cellular source of this immunomodulatory paracrine signaling from the bursa may be due to its resident population of MSCs and/or other immunomodulatory cells. The bursa MSC cell population has been widely characterized from pathological human tissues, and its presence in the native and activated bursa was supported by the results here (*19*). These findings of a beneficial role of the bursa in rotator cuff pathology and healing deviate from the current paradigm in the literature that regards the inflamed bursa as detrimental to rotator cuff health.

Following resolution of the inflammatory phase of healing, the bursa maintained longer-term native tissue properties in the intact rotator cuff that were lost when the bursa was removed. This demonstrates that the bursa protects intact tendons from tissue degradation by inflammatory factors from a nearby tendon injury. Protection against degradation is also supported by the gene expression results at day 56, wherein expression of known tendon-ECM components COL1 and COL3 were increased by the bursa and decreased in its absence. The resulting loss of mechanical integrity in the absence of the bursa is striking and should be considered carefully in the context of the common clinical practice of bursectomy. It is possible that progression of rotator cuff pathology into the healthy cuff may be prevented or delayed by retaining the bursa. Furthermore, cortical and trabecular properties in the humeral head were also regulated by the paracrine environment maintained by the bursa. In this model, the loading environment was the same between both shoulders (i.e., both shoulders had one injured cuff tendon with the rest intact). Therefore, changes in the underlying bone were a result of the tendon-to-bone healing environment, and not the loading environment, and bursa-dependent changes to humeral head bone morphometry likely resulted from paracrine regulation maintained by the bursa. Interestingly, across most cortical bone measurements and trabecular bone volume, the bursa maintained near-native properties that deviated from normal when the bursa was removed. Together, these findings in the intact rotator cuff and underlying humeral bone are evidence against the clinical use of bursectomy, given the implications of the procedure on further rotator cuff pathogenesis.

Having established a role of the bursa in regulating the paracrine environment in the shoulder to protect the uninjured tissues from degradation, the current work then examined the role of the bursa in regulating tendon healing in the injured supraspinatus tendon. Unlike the intact infraspinatus tendon, the bursa increased inflammatory gene expression and cellular infiltration in the healing supraspinatus tendon. Inflammation has only recently begun to be appreciated for its importance in tendon healing after decades of conflicting reports (*26–28*). In particular, the recent literature has emphasized the importance of inflammation *resolution* over *elimination* (*28*). Many of the transcription-level results demonstrated here indicated an upregulated inflammatory response in the presence of the bursa at the earliest timepoints studied. Considering the current understanding that inflammation is an essential first step in the wound healing process, these findings are not definitive on whether the bursa is beneficial (i.e., necessary inflammation activation) or detrimental (i.e., prompting chronic inflammation) in tendon healing. The cell-type transcriptional analysis on the injured supraspinatus tendon indicated an increased presence of cell populations that are important to the healing process, including MSCs, fibroblasts, and vascular cells, in the presence of the bursa. These results suggest that the inflammatory response may incite infiltration of key players that are essential to the healing process. Later in the healing process, the functional consequence of these transcription level changes appeared to be limited, at least as demonstrated by tissue level mechanical properties. Nevertheless, promotion of inflammation by the bursa early in the tendon’s response to injury did not have a detrimental effect on tendon healing. On the other hand, the bursa did promote ECM- and wound healing-related transcription, suggesting that a longer healing period (i.e., greater than 56 days) may permit detection of a functional difference imparted by the bursa on the repaired tendon.

The beneficial role of the bursa in regulating homeostasis in the intact tissues of the shoulder implies that the clinical practice of bursectomy is ill-advised. With a potential shift in practice away from bursectomy, the bursa now presents itself as an ideally situated therapeutic command center for regulating tendon healing. Therefore, the final component of the current study examined the potential to therapeutically target the bursa. Using *in vitro* and *in vivo* model systems, Dex was validated as an attractive drug for prompting the bursa to modulate tendon inflammatory responses. Dex has received recent traction in the literature for inflammation reduction across a number of tissues, including the gingiva, cartilage, and knee synovium, motivating our choice (*29, 30*). The results from *in vivo* testing of this treatment strategy supported the efficacy of Dex delivery to the bursa for decreasing expression of inflammatory cytokines in both the injured and intact rotator cuff tendons. This is consistent with a report that Dex decreased expression of inflammatory genes in the synovium and infrapatellar fat pad and was chondroprotective in a model of post-traumatic osteoarthritis (*29*). This therapeutic approach offers many advantages over directly injecting the tendon, as the fatty nature of the bursa permits retention and delivery of the drug to the bursa, with slow release to the underlying tendon, avoiding detrimental effects of high dose corticosteroids on the tendon. Intraarticular injection of PLGA-based extended release triamcinolone acetonide is FDA-approved for treating osteoarthritis pain in the knee and is being studied for OA pain in the shoulder (*31*). This supports the translatability of microsphere-encapsulated corticosteroids for treating joint pathologies. Future iterations on the candidate drug choice, dose, and timing of injection will refine this bursa-targeted therapeutic approach for clinical translation. Furthermore, the *in vitro* bursa-tendon crosstalk model will allow for high throughput screening of additional candidate drugs for tendons *in vivo*. This approach is similar to that employed in organ-on-a-chip models and will expedite drug screening for clinical translation to improve rotator cuff healing outcomes (*32*).

Studying the bursa through patient biopsies provided important clinical evidence but was limited by *(i)* lack of availability of healthy samples and *(ii)* variability in patient demographics and medical histories for factors known to be associated with rotator cuff disease. Expanding the enrollment of patients in biopsy-focused studies will be a focus of future efforts to develop precision medicine approaches for patient-specific rotator cuff treatment strategies. For the *in vivo* investigations on the role of the bursa in its paracrine environment, emphasis was placed on minimizing total tissue damage and risk of infection by using the smallest skin and muscle incisions possible that allowed for adequate tissue visualization. During the surgical studies, it became apparent that there were inconsistencies in achieving complete bursectomy with the limited size of the trapezius muscle dissection. For the *in vivo* translation of Dex delivery to the bursa, Dex-MS were suspended in sterile saline for injection. Due to the puncture into the bursa for the injection and the volume of the saline bolus it is likely that some Dex-MS escaped the bursa into the subacromial space. Clearance rates of non-immobilized microspheres in the rat shoulder have not been investigated, so it is possible that some of the effect of the bursa treatment strategy was due to direct delivery of Dex to the rotator cuff. However, if this combined approach for therapeutic treatment via redirecting of both the bursa and rotator cuff cellular signaling is effective, it is not necessarily a limiting feature of the delivery strategy.

In conclusion, we observed bursa-tendon crosstalk in rotator cuff tendinopathy in model systems and determined that the bursa regulates the paracrine environment within the shoulder. Specifically, the bursa maintained native properties of the intact rotator cuff and underlying humeral head and regulated the healing response in the injured rotator cuff. Finally, we demonstrated the therapeutic potential for targeting the subacromial bursa with Dex to regulate the inflammatory response in the injured rotator cuff.

## MATERIALS AND METHODS

### Study design

The goal of this study was to investigate the biological role of the subacromial bursa in rotator cuff pathology. We compared proteomic signatures between bursa of patients with a range of rotator cuff pathology diagnoses (subacromial impingement; small, medium, large, massive rotator cuff tears) to determine the presence of clinically relevant bursa-tendon crosstalk. We further explored the role of the bursa in regulating its paracrine environment *in vivo* using the rat rotator cuff injury and/or repair models with or without bursectomy, emphasizing the bursa’s role in both the injured and intact tissues of the shoulder. We investigated bursa-tendon crosstalk in a new *in vitro* bursa explant culture model. Finally, we hypothesized that Dexamethasone delivery to the bursa would regulate the inflammatory response to injury in the rotator cuff. Statistical analysis and samples sizes were determined using power analyses based on the effect size in preliminary experiments. Samples for biomechanical testing, microcomputed tomography (μCT), and histological cell counting were randomized and analyzed by a blinded investigator.

### Study approval

Human study procedures and protocols were approved by the Columbia University Institutional Review Board. Full informed consent was obtained from all patients. Animal studies were approved by the Columbia University Institutional Animal Care and Use Committees.

### Clinical samples

Samples of subacromial bursa tissue were obtained from patients with subacromial impingement or rotator cuff tears undergoing surgery (subacromial decompression or rotator cuff repair/reconstruction, respectively) using a standard arthroscopic technique. The subacromial bursa was biopsied inferior to the acromion; the injured tendon was biopsied at the site of injury. Two to three biopsies of each tissue were obtained per patient. Patient demographics (sex, ethnicity, race, age) and clinical metadata (diagnosis, injury occurrence, steroid history) were prospectively obtained at presurgical office visits.

### Proteomics analysis

Tissues intended for proteomics assessment were stored in AllProtect (Qiagen) and later submitted for mass spectrometry based tandem mass tag (TMT) proteome quantification at the Columbia University proteomics core. The tissues were lysed in urea, digested in Trypsin and Lys-C protease mix, TMT labeled (to allow for multiplexing), fractionated (to increase detection of low-abundance proteins), and run through the mass spectrometer (to quantify the relative abundances of proteins within each tissue sample). The abundance data was the input to all further data analysis using Perseus proteomics software and Enrichr. Patient data (age, sex, ethnicity, race, diagnosis, injury occurrence, steroid use history) and sample designation (bursa subtype) were assigned to each sample prior to analysis. Hierarchical heat map clustering and PCA were performed in Perseus to provide unbiased information on sample similarity via cluster formation. Significantly up and downregulated proteins between groups, as determined by the statistical approach below, were plotted in heats maps by log2 fold change and were the inputs to downstream IPA and Enrichr analyses for gene ontology (GO).

### Acute injury with/without bursectomy

Rats weighing 250-300g (n=3 female, n=3 male for 3-day gene expression; n=2, n=3 for 7-day gene expression; n=3, n=3 for 14-day gene expression) were anesthetized and positioned on their left side. With the right forelimb externally rotated, the skin was palpated to identify the acromial-clavicular bony arch and a skin incision was made 1 cm medial and distal to the arch, exposing the trapezius and the deltoid, respectively. The muscle fascia was removed using blunt dissection. The prominent vein running anterior to posterior across the bony arch was cauterized superficially. An 11-blade was then used to resect the trapezius from the clavicle, acromion, and scapular spine. When the muscle was released, a retractor was placed beneath the trapezius to provide better visualization of the underlying subacromial bursa. Remaining muscle fibers connected to the superior side of the bursa were blunt dissected, and the bursa was grasped with micro-forceps with teeth deep to the trapezius, at its medial margin. Tension was applied to gently pull the bursa away from its loose attachment to the supraspinatus muscle on its inferior side. This process was repeated distally towards the acromion until all the visualized bursa was removed and clear margins were visible where the bursa was previously attached to the acromion. With the bursectomy complete, the deltoid muscle was blunt dissected away from the distal edge of acromion and the opening in the muscle was expanded using an 11-blade until the supraspinatus tendon insertion was visible. Insertion and tendon visualization were improved by holding the forepaw down on the surgical table while adducting the forelimb. The forelimb was internally and externally rotated to identify the margins with the infraspinatus tendon posteriorly and the biceps tendon anteriorly. The supraspinatus tendon was sharply detached from the humeral head and allowed to retract. The deltoid and trapezius were reflected back into their anatomical positions and repaired using 5-0 Prolene suture, and the skin was closed using 4-0 Ethilon suture. The rat was then repositioned onto its right side and the left forelimb was positioned in the same manner. The surgery proceeded identically for the left shoulder, but without a bursectomy. Animals were euthanized at 3, 7, or 14 days after surgery.

### Acute injury and repair with/without bursectomy

Rats weighing 250-300g (n=2 female, n=3 male for 14-day gene expression; n=3, n=2 for 28-day gene expression; n=2, n=3 for 56-day gene expression; n=5, n=6 female for 56-day biomechanical testing) were anesthetized and laid on their left side. The acute injury and repair with bursectomy proceeded in the same manner as described above until the margins of the supraspinatus with the infraspinatus tendon posteriorly and the biceps tendon anteriorly were identified. Next, a 5-0 Prolene suture was used to place a modified Mason-Allen stitch in the supraspinatus tendon. After the tendon grasping suture was passed, the supraspinatus tendon was sharply detached from the humeral head. The insertion site was debrided of any remaining tendon fibers with a burr-tip attachment on a Dremel. A drill-tip attachment on the Dremel was used to pass a bone tunnel in the anterior to posterior direction through the humeral head. Immediately after removing the drill, a 20-gauge needle was passed anterior to posterior through the bone tunnel. The needle on the grasping suture was inserted into the tip of the 20-gauge needle and guided back through the bone tunnel, posterior to anterior, before tying two knots to repair the supraspinatus tendon to its original attachment site. The deltoid and trapezius were subsequently repaired, and skin was closed as described above. The rat was repositioned onto its right side and the acute injury with repair and no bursectomy was performed on the left shoulder. Animals were euthanized 14, 28, or 56 days after surgery.

### Bursa explant dissection, equilibration, and activation

Bursa explants were dissected from rat shoulders according to the bursa dissection protocol (supplementary). These explants were immediately submersed in cold PBS supplemented with 1% pen-strep. Explants were then transferred to bursa complete media containing DMEM/F-12 supplemented with 10% MSC-verified FBS and 1% pen-strep. Explants were equilibrated in this media for 24-48 hours to remove any cells at the tissue margins that died during dissection. To simulate the activated state created in the bursa by a tenotomy, the bursa explants were treated with 20ng/mL IL1β in bursa complete media for 72 hours before co-culture with tendon fibroblasts.

### In vitro model of bursa-tendon crosstalk

Tendinopathy was modelled *in vitro* with IL1b pre-treated tendon fibroblasts (*37*). To investigate crosstalk between the bursa and the tendon, tendon fibroblasts were co-cultured with bursa explants. At the start of co-culture, the media on the tendon fibroblasts (either IL1β-treated or not) was exchanged for 3mL bursa complete media and equilibrated bursa explants (IL1β-activated or untreated control) were added to the wells. The bursa explant and tendon fibroblast monolayer were not in physical contact, as the adipose nature of the bursa caused the bursa to float. The explants and tendon fibroblasts were co-cultured for 24 hours prior to downstream analyses.

### In vitro therapeutically directed bursa-tendon crosstalk

The *in vitro* model of bursa-tendon crosstalk was expanded for use in screening candidate inflammation-regulating drugs for bursa-targeted delivery. Bursa and tendon fibroblasts were activated to an inflamed state for 72 and 24 hours, respectively, with 20ng/mL IL1β as described above. At hours 24 and 48 hours into the 72-hour bursa activation, solubilized Dex (100μM) was added to the culture media of the bursa intended for the activated+Dex group. The same Dex dosing scheme was used for bursa of the Dex only group (without IL1β activation). A range of doses (100nM, 1000nM, 100μM) were confirmed to similarly reduce IL6 and COX2 expression in activated bursa before experiments proceeded (pilot data not shown). RNA was isolated from treated bursa explants to quantify the transcriptional changes resulting from IL1β and Dex treatments.

Following the respective activation protocols for bursa explants and tendon fibroblasts, treated or untreated bursa explants were transferred into co-culture with tendon fibroblasts in fresh bursa complete media. The bursa and tendon fibroblasts were co-cultured for 24 hours prior to downstream analyses.

### Dex-MS bursa-targeted delivery

The efficacy of Dex-MS delivery to the bursa for regulating the inflammatory response in the rotator cuff tendons was established using tendinopathy induced by bilateral supraspinatus tenotomies without bursectomy. Dex-MS (6ug Dex) suspended in 3uL sterile saline bolus was injected into right-side bursa; an equal mass of unloaded MS in a 3uL sterile saline bolus was injected into left-side bursa. Delivery of MS was performed either immediately following supraspinatus tendon injury (concurrent) or 3 days following tendon injury (delayed). The concurrent study was assessed for gene expression changes in the bursa, supraspinatus tendon, and infraspinatus tendon 3 days post-intervention. For the delayed study, supraspinatus tenotomies were performed without drug delivery to the bursa. After 3 days, skin was reopened, the trapezius muscle was reflected and MS (right: Dex-MS; left: unloaded MS) were injected into the bursa. After 4 days (7 days following tendon injury), the bursa, supraspinatus tendon, and infraspinatus tendon were assessed for gene expression changes.

### Statistical analysis

For the clinical proteomics analysis, protein abundances were normalized using Z-scores and multiple sample t tests with a false discovery rate of 0.05 were conducted to identify those proteins that were significantly different between tissue categorizations.

Transcriptomics data from rat bursa and supraspinatus tendon were analyzed using DESeq2 with statistical significance (p<0.05) determined using the adjusted p-value.

All RT-qPCR gene expression fold changes were calculated using 2^-ΔΔCT^ and plotted as log_2_ fold change. Gene expression changes in the tenotomy activated bursa were compared to control bursa using unpaired t-tests with Welch’s correction. Supraspinatus and infraspinatus tendon groups BI, BR, and CTL gene expression changes were compared using ANOVA with Tukey correction for multiple comparisons. Paired right (BR) and left (BI) supraspinatus and infraspinatus groups were also compared using paired student’s t-tests between groups. Statistical significance was set to p<0.05.

Similarly, paired right (BR) and left (BI) supraspinatus and infraspinatus biomechanical testing results and paired right and left humeral head morphometry results were compared using paired student’s t-test between groups. BI and BR groups were compared to CTL for biomechanics and bone morphometry using ANOVA with Tukey correction for multiple comparisons.

Statistical significance for gene expression and NO release changes in the *in vitro* bursa-tendon crosstalk with or without bursa-targeted Dex treatment was determined using one-way ANOVA with a Tukey correction for multiple comparisons and paired student’s t-tests between the paired explants. Statistical significance for gene expression changes in the *in vivo* acute injury model with Dex-MS delivery to the bursa was determined using paired student’s t-tests comparing Dex-MS to unloaded MS control.

## List of Supplementary Materials

Supplementary methods

Figure S1

Figure S2

Figure S3

Figure S4

Figure S5

Figure S6

Figure S7

Figure S8

Data file: proteomics

Datafile: transcriptomics

## Acknowledgements

The authors acknowledge Yelena Akelina of the Microsurgery Research and Training Laboratory for providing rat carcasses for model development. The authors also acknowledge the Proteomics Core and the Genome Center at the Columbia University Irving Medical Center for performing the proteomics and transcriptomics assays, respectively. The authors finally acknowledge the Histology services from the Molecular Pathology core for all histological processing and tissue staining.

## Funding

This study was funded by the National Institutes of Health (R01 AR057836 to ST and K08AR072092 to DK) and National Science Foundation (DGE–1644869 to B.P.M.).

## Author contributions

Conceptualization: BPM, ST

Methodology: BPM, KGB, LS, AJL1, AJL2, CTH, ST

Investigation: BPM, XER, JAK, ACI, AJL1, LAF

Visualization: BPM, XER, JAK, AJL2

Funding acquisition: ST, DK

Project administration: ST

Supervision: ST

Writing – original draft: BPM, ST

Writing – review & editing: XER, JAK, ACI, AJL1, KGB, AJL2, LS, DK, CTH, WNL, ST

## Competing interests

The authors declare that they have no competing interests.

## Data materials availability

All data associated with the study are present in this paper, in the Supplementary Materials, or submitted to a data repository (human bursa and tendon proteomics, rat bursa and tendon transcriptomics).

## Notes

### Competing Interest Statement

The authors have declared no competing interest.

