## Supplemental Materials for "The subacromial bursa is a key regulator of the rotator cuff and a new therapeutic target for improving repair"

**Supplementary Materials:**

#### SUPPLEMENTARY METHODS

##### Gene expression

At the post-operative timepoints detailed in the study design, rats were sacrificed and the bursa, supraspinatus, and/or infraspinatus were dissected and stored in Allprotect (Qiagen) at room temperature for 24 hours and then stored at -80 °C. Before RNA isolation, the tissue was placed within a nested stack of Eppendorf tubes, the smaller of which had a hole in its bottom, and centrifuged at 500g for 30 seconds to remove excess Allprotect. Explants previously dissected for the *in vitro* tendinopathy model were removed from culture and snap-frozen without Allprotect. Snap-frozen issues were then homogenized using a ball mill homogenizer (Mikro-Dismembrator U, Sartorius). RNA extraction was conducted using QIAzol Lysis Reagent (Qiagen) and MaXtract High Density phase lock gel tubes (Qiagen). RNA isolated from bursa-derived cells undergoing the multilineage differentiation assay or tendon fibroblasts from the *in vitro* bursa-tendon crosstalk model did not undergo the above steps of preparation but were instead treated with the RNeasy Mini Kit (Qiagen) RLT buffer to produce a cell lysate. RNA purification was conducted using RNeasy Mini Kit spin columns (Qiagen) according to manufacturer protocol. Purified RNA was quantified and checked for quality using a spectrophotometer (Nanodrop 1000, Thermofisher Scientific). RNA samples intended for RNA sequencing were further quantified and checked for quality using the Bioanalyzer (Agilent) through the Columbia University Molecular Pathology core service. RNA samples intended for quantitative real-time polymerase chain reaction (RT-qPCR) were reverse transcribed to cDNA using the High-Capacity cDNA Reverse Transcription kit (Applied Biosystems). RT-qPCR was conducted in the QuantStudio 6 Flex (Applied Biosystems) using PowerUP SYBR Green Master Mix (Applied Biosystems). Gene expression changes were measured for inflammatory (COX2, IL1b, IL6, IL10, TGFβ, SDF1a, TNFα) and tenogenic/chondrogenic/ECM (SCX, SOX9, RUNX2, COL1, COL3, BGN) genes in the inflammatory (bursa, infraspinatus, supraspinatus) and reparative (infraspinatus and supraspinatus) healing phases, respectively.

##### Microcomputed tomography

For rat whole-shoulder anatomical scans, samples that included the humerus, scapula, and attached soft tissues were dissected. Samples were fixed in 10% formalin in 1x PBS for 1 day and subsequently stained in 0.276 M $HgCl_2$ in $dH_2O$ for 1-2 days *(34)*. Samples were rinsed, mounted, and scanned in air (Skyscan 1272, Aartselaar, Belgium). Kimwipes soaked in PBS were used to line the containers to prevent drying during scanning. Samples were scanned at an energy of 57 kVP, Al 0.25 filter, and resolution of 19.1 μm and reconstructed (nRecon, Bruker). Images were visualized using DataViewer (Bruker).

Supraspinatus-infraspinatus-humerus units were dissected from each side of euthanized rats 56 days after supraspinatus injury and repair with or without bursectomy for microcomputed tomography (μCT) bone morphometry analysis (n = 11). Distal ends of the tendon-humerus units were embedded in agarose so that the tendons hung loosely in air in line with the scanning axis. Samples were scanned at an x-ray tube potential of 50 kVP, intensity of 180 uA, 1900 ms exposure time, with an Al 0.25mm filter, and isometric resolution of 5 μm (Skyscan 1272, Bruker). The scan data were reconstructed with the provided CT software (nRecon, Bruker) using alignment optimization and beam-hardening correction. Reconstructed images were visualized with compatible programs (DataViewer and CTVox, Bruker). Two hydroxyapatite calibration phantoms were used to relate CT values to a mineral-equivalent value for cortical and trabecular bone morphometry. Reconstructed images were evaluated using a segmentation algorithm to separate cortical and trabecular bone of the humeral head proximal to the growth plate (CTAn, Bruker). The morphometric measurements included periosteal envelope cross-sectional area, cortical measurements (bone area, area fraction, cross-sectional thickness), and trabecular measurements (bone volume, volume fraction, number, thickness, separation).

**Tendon-to-bone biomechanics**

After microcomputed tomography scans were complete, supraspinatus-infraspinatus-humerus units from the 56-day timepoint were biomechanically tested. All samples were tested in a saline bath at 25°C to prevent thermal collagen denaturation on a table-top tensile tester (Electroforce 3230, TA Instruments) fitted with a 100-lb load cell (TA instruments). Before testing, the supraspinatus muscle was removed from tendon. Samples were placed into custom 3D-printed fixtures that secured samples in uniaxial tension *(35)*. Supraspinatus tendons were secured between Kimwipes with a drop of cyanoacrylate adhesive (Loctite, Ultra Gel Control) before mounting onto custom grips. For all mechanical testing protocols, samples were first pre-loaded to 0.05 N, pre-conditioned by applying 5 cycles of sinusoidal wave consisting of 5% strain and 0.2%/s, and rested for 300 seconds. Samples were then strained in tension at 0.2%/s to failure. Infraspinatus tendon mechanical testing was performed after supraspinatus testing was completed. Before testing the infraspinatus tendon, the infraspinatus muscle was carefully removed from the tendon. Samples were placed into custom 3D-printed fixtures, unique from those used for supraspinatus testing, that oriented the infraspinatus tendon in uniaxial tension.

Enthesis structural properties, such as failure load and stiffness were determined from load-deformation curves. Stiffness was calculated by a MATLAB (Matlab2019a, MathWorks) custom algorithm that identifies the best fitting line within a sufficient bin width (i.e., remove data below 10% of max load and above 95% of max load) by implementing the random sample correlation (RANSAC) technique *(36)*.

**Transcriptomics analysis**

RNA isolated from the subacromial bursa and supraspinatus tendons from the 3- and 14-day post-operative timepoints were analyzed using RNA sequencing (JP Sulzbger Columbia Genome Center). Briefly, total RNA, isolated according to the above protocol, went through poly-A pulldown mRNA purification and was reverse transcribed to cDNA. The cDNA was broken into a library of small fragments, oligonucleotide adapters were attached, and samples were sequenced. The reads from the sequencing were aligned to a reference genome. The aligned genome was analyzed for significant differences in expression of transcripts using DESeq2 (adjusted p <0.05 significant).

Gene lists were sorted by significance based on the DESeq2 analysis (adjusted p-value <0.05 considered significant). Significantly differentially expressed genes were separated into lists of up- and down-regulated genes, relative to control, based on their log fold change (positive: upregulated; negative: downregulated). These lists were separately uploaded to Enrichr for computing gene set enrichment in order to identify differentially regulated pathways, GO, and cell-types using built-in algorithms *(33)*.

**Histology**

Clinical tissue samples intended for histological analysis were fixed in 10% formalin for 24 hours, rinsed and stored in 70% ethanol, and embedded in paraffin blocks. Seven micrometer tissue sections were stained with Masson's Trichrome to depict differences in tissue morphology. Whole-slide scanning images were taken at 20x.

Rat whole shoulders or subacromial bursa tissues were dissected, fixed in 10% phosphate-buffered formalin for 24 hours, decalcified in formic acid for 14-21 days (only for samples containing bone), and stored in 70% ethanol before paraffin embedding. Whole shoulders were oriented and trimmed prior to embedding such that coronal sections containing supraspinatus tendon, bursa, and acromion were collected upon further sectioning. Sections (7 μm) were then obtained from paraffin blocks and stained with hematoxylin and eosin (H&E) or Movat's pentachrome. Brightfield images were taken at 5x and 20x (for cell quantification) magnification.

**Tendon fibroblasts isolation, culture, and activation**

Supraspinatus and infraspinatus tendons from the rat rotator cuff were dissected from the shoulder of control rats. All visible muscle was removed from the tendon using a #15 scalpel blade. The tendons were added to 4mg/mL Collagenase Type-II in 5mL of serum-free MEMα treated with 1% penicillin-streptomycin (pen-strep) in a 50mL Falcon tube. The tubes were placed on an orbital shaker at 190rpm and 37°C and were checked every 30 minutes until the media was opaque without remaining tissue pieces. This solution was filtered through a 70-100mm mesh cell strainer over a clean 50mL Falcon tube. The cell suspension was pelleted by centrifugation at 500g for 5 minutes. The collagenase-containing media was removed, and the cells were resuspended in tendon fibroblast complete media containing MEMα, 10% fetal bovine serum (FBS), and 1% pen-strep. These cells were pelleted once more and resuspended in complete media before plating. Media was changed every 2-3 days; cells were passaged at 80% confluence and used at passage 2-4.

For experiments with IL1β pre-treated tendon fibroblasts, passage 2-4 tendon fibroblasts were lifted and plated at a density of 40,000 cells/well in tendon fibroblast complete media in a 12-well tissue culture-treated plate. After attaching to the plate for 24 hours, 20ng/mL IL1β was added to tendon fibroblast complete media and kept on the cells for 24 hours before co-culture. For experiments without IL1β pre-treatment, this IL1β treatment was omitted.

**Cellularity measurements**

Using ImageJ software (NIH), hematoxylin (nuclear) and eosin (cytoplasmic/ECM) stained whole shoulder histological sections were converted to 8-bit, made binary, and the cell numbers were determined using the "analyze particles" function within an ROI containing the tendon from the border between scar and humeral head to just prior to musculotendinous junction. Sections were blinded to experimental group prior to analysis.

**Bursa dissection**

Rats were euthanized via carbon dioxide asphyxiation, soaked in ethanol, and laid on the side on the dissection table. While externally rotating the forelimb, the arm was palpated to locate the acromioclavicular arch beneath the skin. When located, a 4 cm lateral incision was made with a 15-blade from 2 cm medial to 2 cm distal to the bony arch. Blunt dissection with scissors through the fascia layer followed and was considered complete when there was no remaining fascia over the deltoid and the trapezius. Next, an 11-blade was used to sharply detach the deltoid on the distal side and the trapezius on the medial side of the acromioclavicular arch, while avoiding rupture of the prominent vein that passes in the anterior to posterior direction over the bony arch. With these muscles detached, the trapezius was gently resected further, taking care to not disturb the underlying subacromial bursa, until the entirety of the bursa was visualized. The bursa was then grasped with a fine tip forceps at its most medial margin, and gently lifted while passing an 11-blade beneath the bursa and above the underlying supraspinatus muscle- separating the two. This maneuver was used to dissect the bursa distally to its connection to the acromion. Here, the 11-blade was angled such that it passed beneath the acromion, separating the bursa from the underside of the acromion. The dissected subacromial bursa tissue was then used for downstream experimentation or analysis.

**Bursa-derived mesenchymal stem cell (BSC) isolation and culture**

Subacromial bursa tissue was dissected from euthanized rats, as described above, and rinsed in cold phosphate buffered saline (PBS) supplemented with 1% penicillin-streptomycin (pen-strep). The tissue was digested in collagenase type II (3mg/mL; Worthington) in complete media (DMEM/F-12 (Gibco) supplemented with 10% fetal bovine serum (Gibco) and 1% pen-strep for 1-2 hours. Any remaining tissue was mechanically disrupted via trituration in an 18G needle and 5mL syringe and then passed through a 100μm cell strainer before pelleting by centrifugation at 350-450g for 5 minutes. The isolated cells were resuspended in complete media and pelleted once more to remove any remnant collagenase. After the second centrifugation, the cells were resuspended and plated in complete media and cultured at 37°C and 5% CO2. Cells were allowed to attach for 48 hours before the first media change. Beyond the first media change, media was replaced every 3 days until the cells reached 60-80% confluence when they were trypsinized for passaging; cells were used between passage 3 to 5.

**Multilineage differentiation**

Induction: BSCs were lifted, counted, and plated according to manufacturer protocols for osteogenic (STEMPRO Osteogenesis Differentiation Kit, Gibco) and adipogenic (STEMPRO adipogenesis Differentiation Kit, Gibco) differentiation. BSCs were lifted, counted, and pelleted according to the manufacturer protocol for chondrogenic (hMSC Chondrogenic Differentiation Medium BulletKit supplemented with TGF-b3, Lonza) differentiation. BSCs were lifted, counted, plated at 1x104 cells/cm2 and induced to tenogenic differentiation in alpha-MEM (Gibco) supplemented with 2% FBS (Gemini), 1% pen-strep, and 100ng/mL BMP-12 for 7-14 days *(38)*.

Gene expression: After 14 days in differentiation medium, RNA was isolated, quantified, and reverse transcribed (as described below) for all samples intended for gene expression. Gene expression changes were measured for the following: osteogenesis (ALP), adipogenesis (LPL, ADIPOQ, PPARgamma), chondrogenesis (ACAN, COL2A1, SOX9), and tenogenesis (SCX, TNMD).

Matrix staining: After 14 days in differentiation medium, BSCs induced towards adipogenesis were fixed in 4% formaldehyde and stained for lipid deposition (HCS LipidTOX Green Neutral Lipid Stain counterstained with DAPI, Invitrogen). Fluorescent images were taken at 5x magnification. After 21 days in differentiation medium, BSCs induced towards osteogenesis were fixed in 10% formalin and stained for calcium deposition (Alizarin Red staining solution, Millipore Sigma). The BSC pellets induced towards chondrogenesis were fixed, embedded in paraffin, and sectioned. These sections (7 μm) were stained for glycosaminoglycans deposition using Alcian Blue stain. Brightfield images were taken at 5x magnification.

**Immunocytochemistry**

BSCs were lifted, counted, plated on human fibronectin-coated glass slides (25,000 cells / 24-well plate well), and cultured overnight. The following day, the cells were formalin-fixed, stained with MSC-defining cell markers anti-CD73 (1:200; Abcam ab175396) and anti-CD90 (1:200; Abcam ab225), incubated for 1 hour with goat-anti-rabbit IgG H&L AF488 (1:500; Abcam ab150081) and goat-anti-mouse IgG H&L AF594 (1:200; Abcam ab150120), and mounted onto microscope slides with anti-fade media containing DAPI *(39)*. Fluorescence images were taken at 10x magnification.

**Flow cytometry**

BSCs were lifted, counted, and stained with conjugated antibodies for MSC-defining cell markers anti-CD90 PE (1:200; Biolegend 205903), anti-CD29 FITC (1:100; Biolegend 102205), and hematopoietic cell marker anti-CD45 APC/Cy7 (1:100; Biolegend 202216). The stained cells were fixed in 1% formaldehyde in cell staining buffer and analyzed using flow cytometry (LSRII Flow Cytometer, Columbia Center for Translational Immunology).

**Live-dead imaging**

Viability of the bursa explants in culture was validated using the Live/Dead Cell Imaging Kit (Invitrogen). Healthy bursa explants were equilibrated for 8 days to validate that the bursa stayed alive beyond the length of the co-culture experiment. Bursa activated with IL1β for 72 hours were also assessed for viability. Bursa treated with 1% Triton-X were used as a kill control comparison.

**Nitric Oxide (NO) release**

At the conclusion of the *in vitro* bursa activation and the *in vitro* crosstalk experiments, media from all wells and fresh bursa complete media were stored and frozen for media-based analysis. Nitric oxide, an indicator of inflammation, released into the culture media by the tendon fibroblasts and bursa explants was quantified using the Griess Reagent System (Promega) according to manufacturer protocol.

**Dexamethasone-loaded microsphere fabrication and characterization**

Dexamethasone 21-phosphate disodium salt (Dex, Sigma D1159) was encapsulated in Poly(D,L-lactide-co-glycolide) (PLGA, 75:25 lactide:glycolide, Sigma P1941) as previously published *(40–42)*. A solution of 10 mg Dex dissolved in 0.5 mL ddH2O was added to a PLGA solution consisting of 200 mg PLGA dissolved in 4.5 mL dichloromethane. The polymer-drug mixture was added dropwise to a 1% poly(vinyl alcohol) solution and stirred at 450 RPM for 4 hours. Unloaded microspheres were fabricated by adding 0.5 mL ddH2O to the PLGA solution. Microspheres were collected and purified by multiple rounds of washing in ddH2O and centrifugation at 1500 RCF and subsequently lyophilized for 48 hours and stored at -20°C until further use.

Microspheres were sputter coated and visualized by using a ZEISS Sigma VP scanning electron microscope. Drug release from Dexamethasone-loaded microspheres (Dex-MS) was determined by suspending 5 mg Dex-MS in 1mL PBS. At each timepoint, 500 µL aliquots were removed from the suspension solution and replaced with an equal volume of fresh PBS. Released Dex was quantified against a standard curve by measuring absorbance at 242 nm (BioTek Gen5).

**Supplemental Figures:**

**Fig. S1: The bursa is activated in patients with rotator cuff tendinopathy.** (A) A breakdown of patient demographics and (B) clinical data revealed variability among the patient population. (C) The bursa proteome clustered by age and tissue subtype. (D) The torn rotator cuff tendon proteome clustered according to tear size according to PCA and (E) hierarchical clustering heat map. (F) Venn diagram of differentially expressed proteins between subtypes revealed 34 unique to the fatty vs. fibrous comparison, 15 unique to the vascular vs. fibrous comparison, and one shared protein (SOD3). (G) Pathway analysis of the bursa proteome revealed biological processes upregulated in the vascular compared to fibrous bursa subtypes, with (H) the heat map of differentially regulated genes indicating specific genes implicated in these pathways.

**Fig. S2: The anatomy of the rat shoulder clinically relevant for investigations related to the subacromial bursa.** (A) Schematic representation of the shoulder indicating the soft and hard tissue landmarks in sagittal and transverse views. (B) Surgical dissection of the shoulder revealed distinct tissue planes, including the bursa inferior to the trapezius muscle and acromioclavicular (A-C) arch and superior to the supraspinatus tendon. (C) Contrast-enhanced μCT of the shoulder revealed the bursa as a region of lower radiopacity between the acromion and supraspinatus tendon.

**Fig. S3: The bursa contains a resident mesenchymal stem cell population.** (A-C) Flow cytometry on bursa-derived cells revealed CD29+, CD90+, CD105+, and (C) CD45- cells. (D) immunocytochemistry on a bursa-derived cell monolayer supported flow cytometry results, indicating positive staining for CD73 and CD90. (E) Cell matrix stained positive for lipid deposition following adipogenic differentiation. (F) Cell matrix stained positive for mineral deposition following osteogenic differentiation. (G) Expression of markers of multilineage differentiations supported the successful differentiation of bursa-derived mesenchymal stem cells into adipogenic, chondrogenic, osteogenic, and tenogenic lineages according to the respective differentiation conditions.

**Fig. S4: The bursa is activated by the inflammatory milieu present in models of tendinopathy.** (A-B) Venn diagrams on genes up- and down-regulated in the activated bursa at day 3 and 14 revealed thousands of shared and unique genes differentially expressed at both timepoints in both directions. (C) Transcriptomic cell-type analysis at days 3 and 14 revealed immune and vascular cells in the activated bursa. (D) Pathway analysis of day 14 bursa revealed specific pathways upregulated in the activated bursa. (E) Gene expression by bursa cells revealed an early increase and later decrease in the activated state of the bursa by day 14.

**Fig. S5: Bursa explants remained viable in culture.** (A) Bursa explants cultured in complete media for 8 days remained viable, as did (B) bursa activated by IL1β for 3 days (LIVE/DEAD stain, green). (C) Kill control induced with Triton-X was used as a comparison for dead cells (LIVE/DEAD stain red).

**Fig. S6: The bursa modulates its paracrine environment.** (A) Gene expression by the intact infraspinatus tendon 14 days after supraspinatus tenotomy revealed no effect of the bursa on inflammatory expression. (B) Co-culture of activated or control bursa with untreated tendon fibroblasts resulted in decreased expression of IL6 and COX2 by the tendon fibroblasts when co-cultured with activated bursa compared to control. (C) Co-culture with activated bursa led to increased secretion of NO into the media. (D) At day 28 following supraspinatus tenotomy and repair, the bursa had no effect on infraspinatus expression of ECM-related genes.

**Fig. S7: The bursa drives transcriptional changes and cellular infiltration in the injured tendon.** (A) Venn diagrams indicate up- and down-regulated genes in the supraspinatus tendon from bursa removed and bursa intact groups compared to control at day 3. (B) COX2 expression was upregulated in the injured supraspinatus tendon by the bursa at day 3. (C) Histological quantification of cellular infiltration into the injured tendon revealed a trending increase in infiltration driven by the bursa ($p=0.099$).

**Fig. S8: Effect of delayed delivery of Dex-MS on injured tendon inflammatory response.** (A) Dex-MS were delivered to the bursa 3 days after supraspinatus tenotomy. (B) Gene expression analysis revealed decreased COX2 ($p=0.071$) and increased SDF1a expression by the injured supraspinatus tendon, (C) increased SDF1 expression by the intact infraspinatus tendon, and (D) no effect on gene expression by the bursa.

### Figure S1

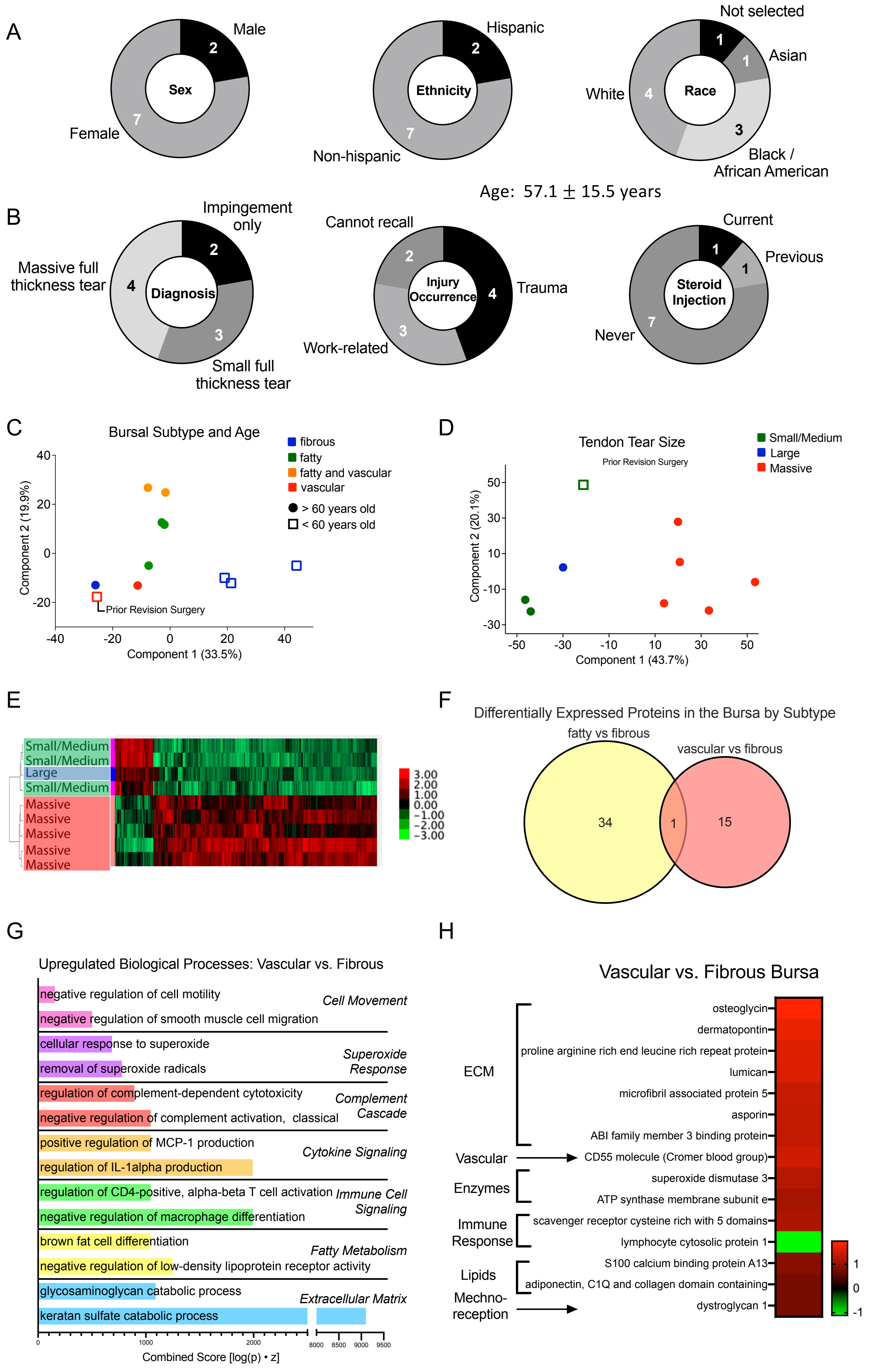

A
Sex
Male
2
Female
7
Ethnicity
Hispanic
2
Non-hispanic
7
Race
Not selected
1
Asian
1
White
4
Black / African American
3
Age: 57.1 ± 15.5 years
B
Diagnosis
Impingement only
2
Massive full thickness tear
4
Small full thickness tear
3
Injury Occurrence
Cannot recall
2
Trauma
4
Work-related
3
Steroid Injection
Current
1
Previous
1
Never
7
C
Bursal Subtype and Age
fibrous
fatty
fatty and vascular
vascular
> 60 years old
< 60 years old
Prior Revision Surgery
Component 2 (19.9%)
Component 1 (33.5%)
D
Tendon Tear Size
Prior Revision Surgery
Small/Medium
Large
Massive
Component 2 (20.1%)
Component 1 (43.7%)
E
Small/Medium
Small/Medium
Large
Small/Medium
Massive
Massive
Massive
Massive
Massive
3.00
2.00
1.00
0.00
−1.00
−2.00
−3.00
F
Differentially Expressed Proteins in the Bursa by Subtype
fatty vs fibrous
vascular vs fibrous
34
1
15
G
Upregulated Biological Processes: Vascular vs. Fibrous
negative regulation of cell motility
negative regulation of smooth muscle cell migration
Cell Movement
cellular response to superoxide
removal of superoxide radicals
Superoxide Response
regulation of complement-dependent cytotoxicity
negative regulation of complement activation, classical
Complement Cascade
positive regulation of MCP-1 production
regulation of IL-1alpha production
Cytokine Signaling
regulation of CD4-positive, alpha-beta T cell activation
negative regulation of macrophage differentiation
Immune Cell Signaling
brown fat cell differentiation
negative regulation of low-density lipoprotein receptor activity
Fatty Metabolism
glycosaminoglycan catabolic process
keratan sulfate catabolic process
Extracellular Matrix
Combined Score [log(p) • z]
H
Vascular vs. Fibrous Bursa
ECM
osteoglycin
dermatopontin
proline arginine rich end leucine rich repeat protein
lumican
microfibril associated protein 5
asporin
ABI family member 3 binding protein
Vascular
CD55 molecule (Cromer blood group)
Enzymes
superoxide dismutase 3
ATP synthase membrane subunit e
Immune Response
scavenger receptor cysteine rich with 5 domains
lymphocyte cytosolic protein 1
Lipids
S100 calcium binding protein A13
adiponectin, C1Q and collagen domain containing
Mechno-reception
dystroglycan 1
1
0
-1

Figure S2

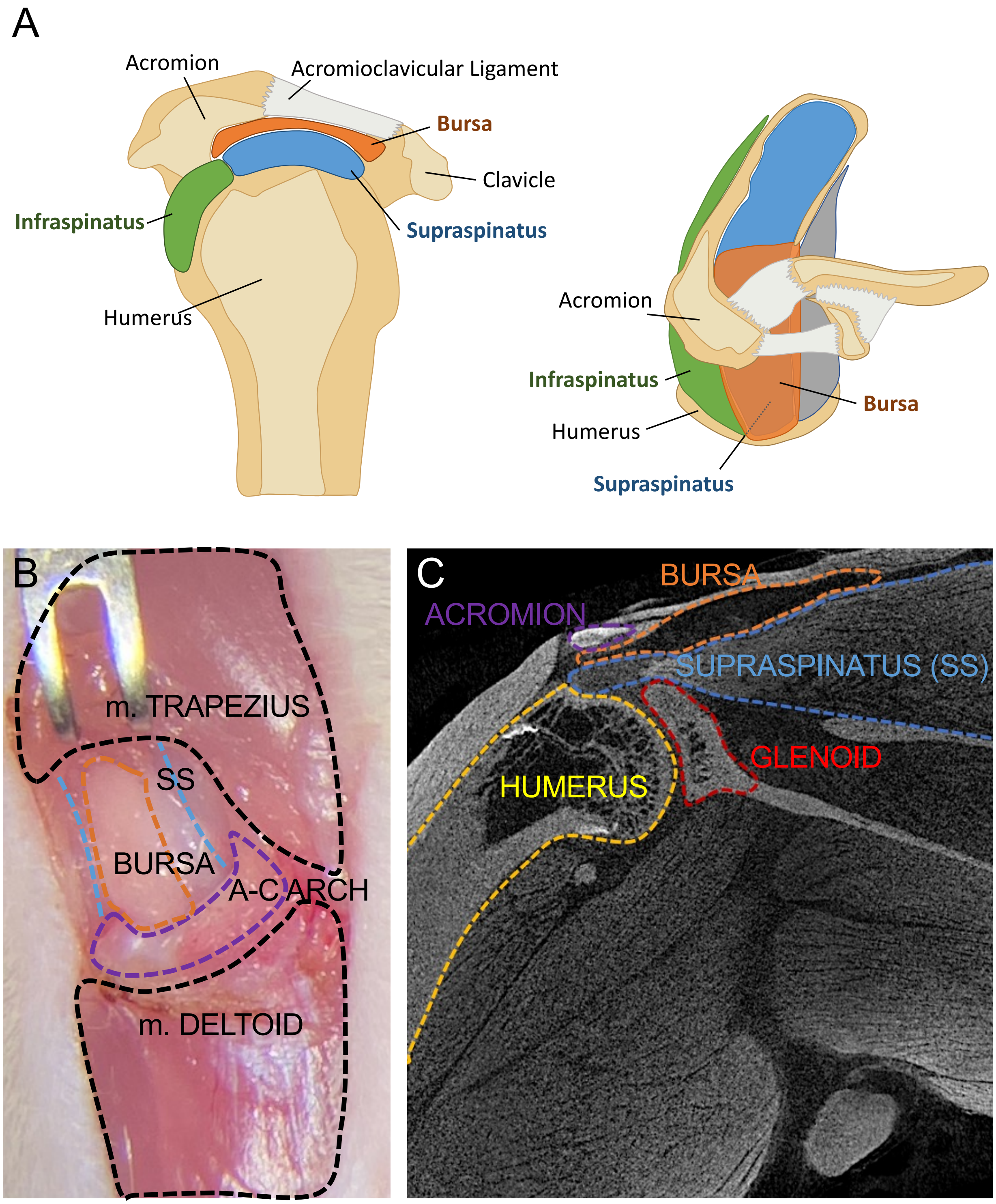
A
Acromion
Acromioclavicular Ligament
Bursa
Clavicle
Infraspinatus
Supraspinatus
Humerus
Acromion
Infraspinatus
Humerus
Bursa
Supraspinatus
B
m. TRAPEZIUS
SS
BURSA
A-C ARCH
m. DELTOID
C
BURSA
ACROMION
SUPRASPINATUS (SS)
GLENOID
HUMERUS

Figure S3

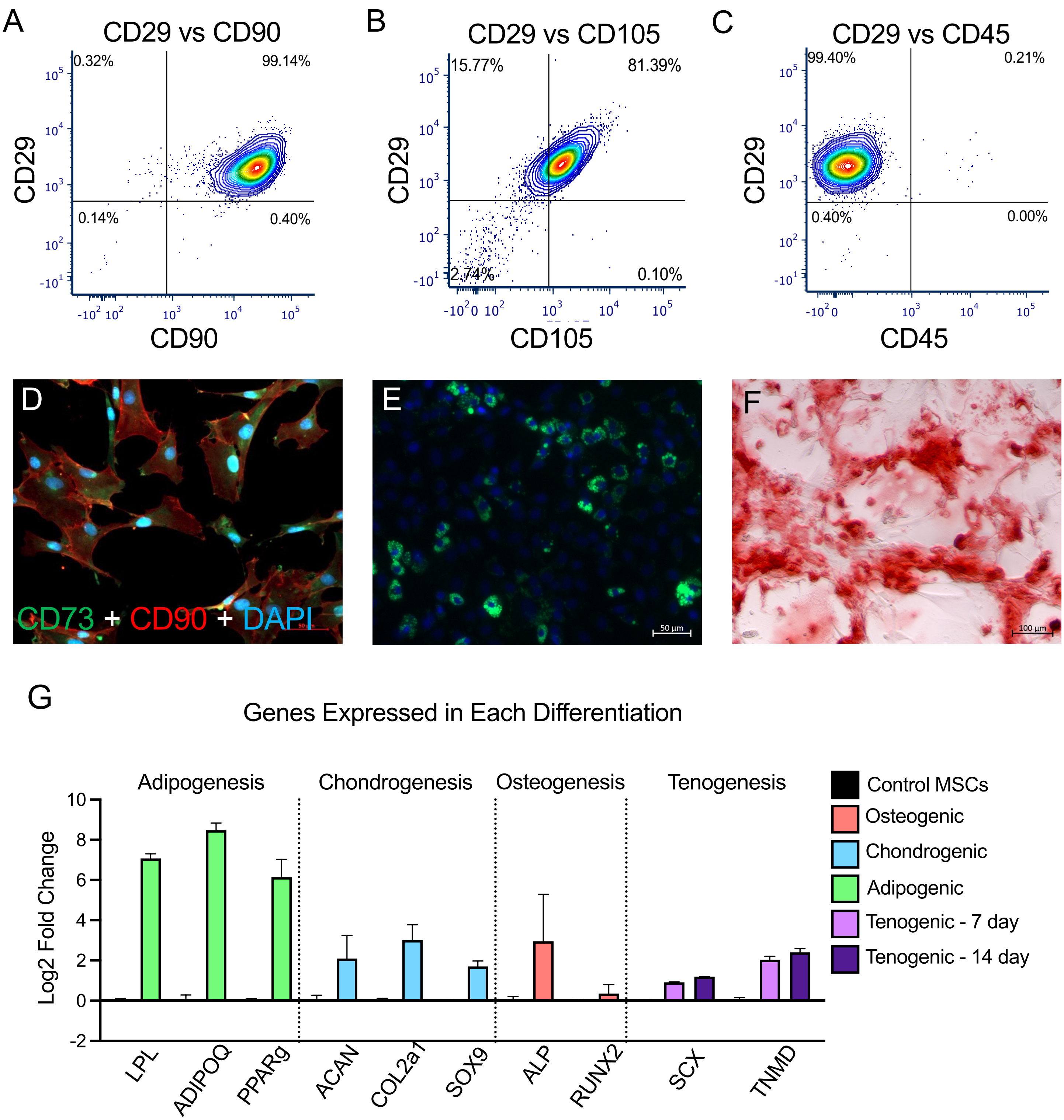
A
CD29 vs CD90
0.32%
99.14%
0.14%
0.40%
CD29
CD90
B
CD29 vs CD105
15.77%
81.39%
2.74%
0.10%
CD29
CD105
C
CD29 vs CD45
99.40%
0.21%
0.40%
0.00%
CD29
CD45
D
CD73 + CD90 + DAPI
E
50 µm
F
100 µm
G
Genes Expressed in Each Differentiation
Adipogenesis
Chondrogenesis
Osteogenesis
Tenogenesis
Log2 Fold Change
LPL
ADIPOQ
PPARg
ACAN
COL2a1
SOX9
ALP
RUNX2
SCX
TNMD
Control MSCs
Osteogenic
Chondrogenic
Adipogenic
Tenogenic - 7 day
Tenogenic - 14 day

### Figure S4

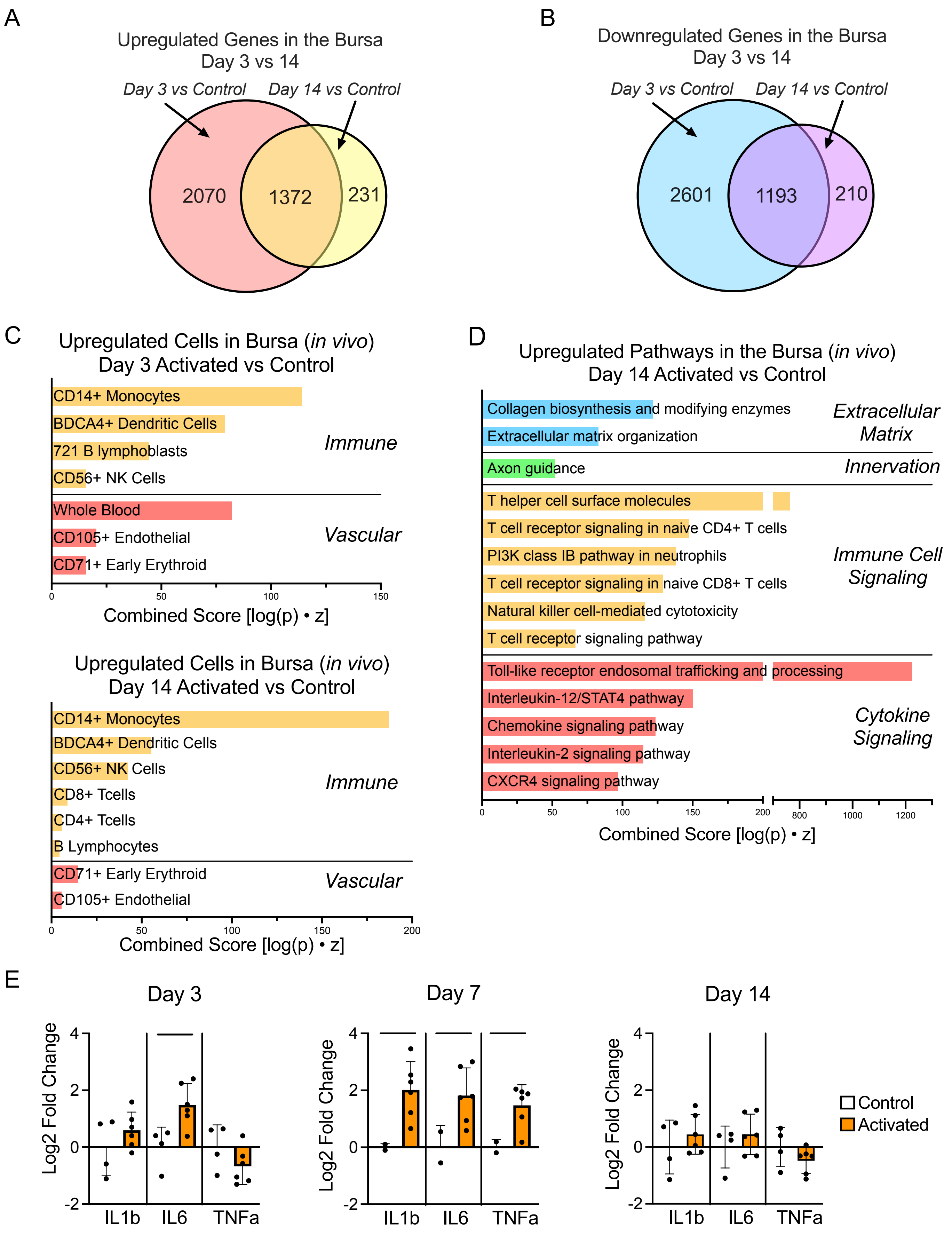
A
Upregulated Genes in the Bursa
Day 3 vs 14
Day 3 vs Control
Day 14 vs Control
2070
1372
231
B
Downregulated Genes in the Bursa
Day 3 vs 14
Day 3 vs Control
Day 14 vs Control
2601
1193
210
C
Upregulated Cells in Bursa (in vivo)
Day 3 Activated vs Control
CD14+ Monocytes
BDCA4+ Dendritic Cells
721 B lymphoblasts
CD56+ NK Cells
Immune
Whole Blood
CD105+ Endothelial
CD71+ Early Erythroid
Vascular
0
50
100
150
Combined Score [log(p) • z]
Upregulated Cells in Bursa (in vivo)
Day 14 Activated vs Control
CD14+ Monocytes
BDCA4+ Dendritic Cells
CD56+ NK Cells
CD8+ Tcells
CD4+ Tcells
B Lymphocytes
Immune
CD71+ Early Erythroid
CD105+ Endothelial
Vascular
0
50
100
150
200
Combined Score [log(p) • z]
D
Upregulated Pathways in the Bursa (in vivo)
Day 14 Activated vs Control
Collagen biosynthesis and modifying enzymes
Extracellular matrix organization
Extracellular Matrix
Axon guidance
Innervation
T helper cell surface molecules
T cell receptor signaling in naive CD4+ T cells
PI3K class IB pathway in neutrophils
T cell receptor signaling in naive CD8+ T cells
Natural killer cell-mediated cytotoxicity
T cell receptor signaling pathway
Immune Cell Signaling
Toll-like receptor endosomal trafficking and processing
Interleukin-12/STAT4 pathway
Chemokine signaling pathway
Interleukin-2 signaling pathway
CXCR4 signaling pathway
Cytokine Signaling
0
50
100
150
200
800
1000
1200
Combined Score [log(p) • z]
E
Day 3
Day 7
Day 14
Log2 Fold Change
4
2
0
-2
IL1b
IL6
TNFa
Control
Activated

### Figure S5

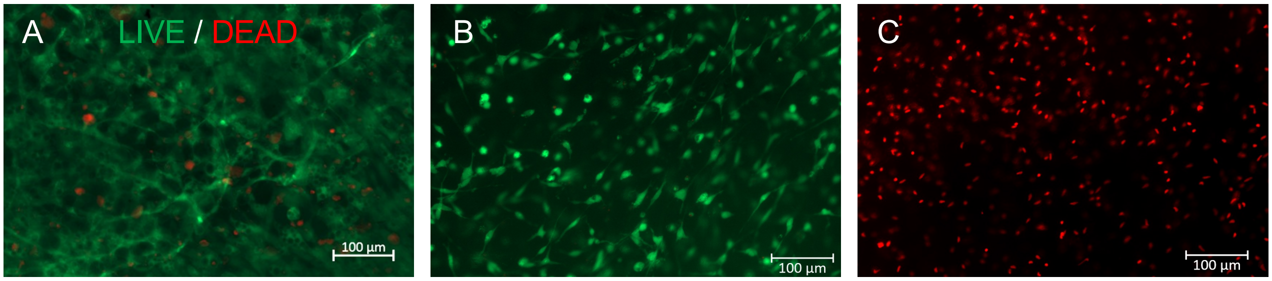

Figure S6

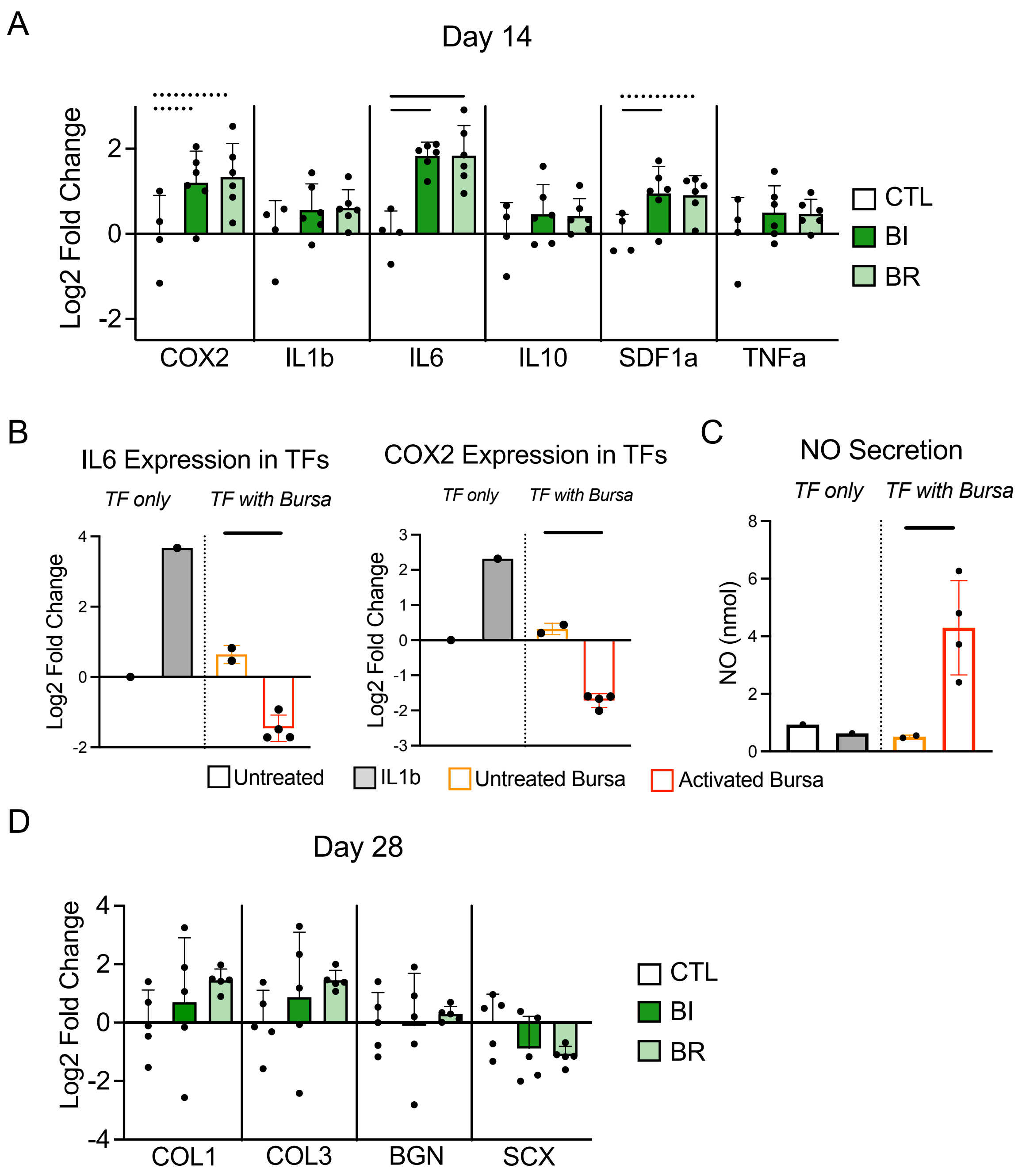
A
Day 14
Log2 Fold Change
2
0
-2
COX2
IL1b
IL6
IL10
SDF1a
TNFa
CTL
BI
BR
B
IL6 Expression in TFs
TF only
TF with Bursa
Log2 Fold Change
COX2 Expression in TFs
TF only
TF with Bursa
Log2 Fold Change
C
NO Secretion
TF only
TF with Bursa
NO (nmol)
Untreated
IL1b
Untreated Bursa
Activated Bursa
D
Day 28
Log2 Fold Change
COL1
COL3
BGN
SCX
CTL
BI
BR

Figure S7

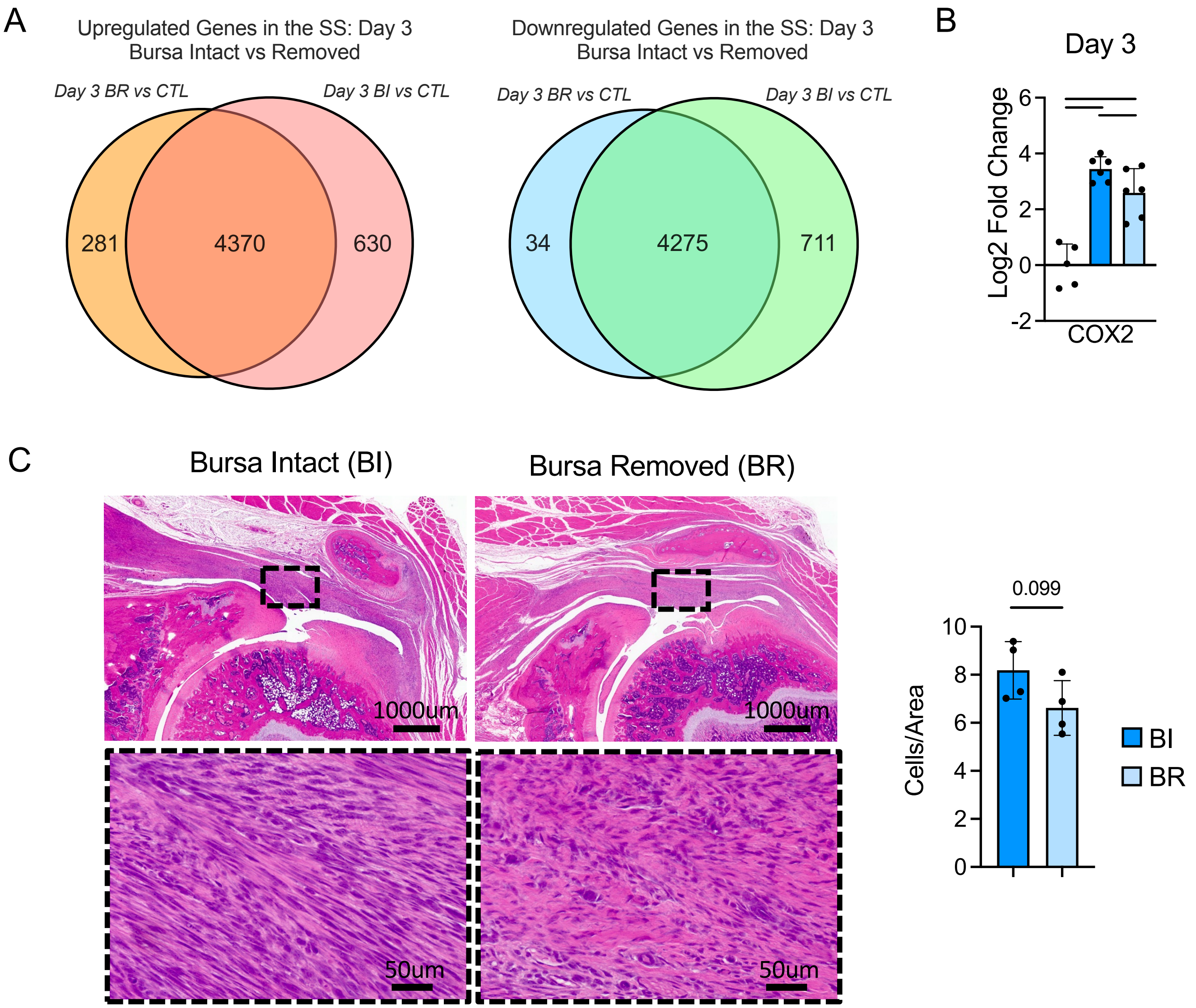
A
Upregulated Genes in the SS: Day 3
Bursa Intact vs Removed
Day 3 BR vs CTL
Day 3 BI vs CTL
281
4370
630
Downregulated Genes in the SS: Day 3
Bursa Intact vs Removed
Day 3 BR vs CTL
Day 3 BI vs CTL
34
4275
711
B
Day 3
Log2 Fold Change
6
4
2
0
-2
COX2
C
Bursa Intact (BI)
Bursa Removed (BR)
1000um
1000um
50um
50um
0.099
Cells/Area
10
8
6
4
2
0
BI
BR

Figure S8

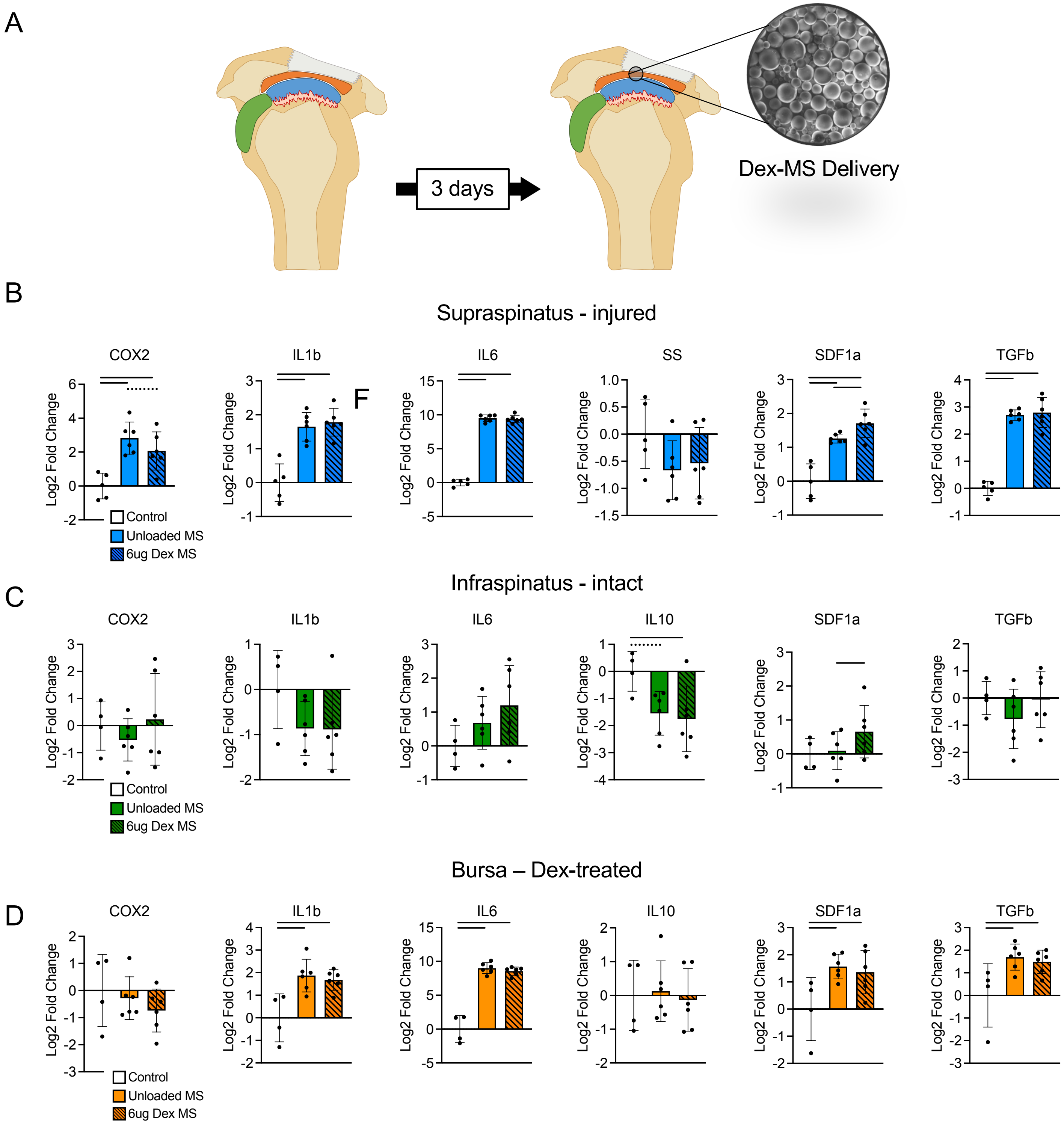
A
3 days
Dex-MS Delivery
B
Supraspinatus - injured
COX2
IL1b
IL6
SS
SDF1a
TGFb
F
Log2 Fold Change
Control
Unloaded MS
6ug Dex MS
C
Infraspinatus - intact
COX2
IL1b
IL6
IL10
SDF1a
TGFb
Log2 Fold Change
Control
Unloaded MS
6ug Dex MS
Bursa – Dex-treated
D
COX2
IL1b
IL6
IL10
SDF1a
TGFb
Log2 Fold Change
Control
Unloaded MS
6ug Dex MS
